# Glioma synapses recruit mechanisms of adaptive plasticity

**DOI:** 10.1101/2021.11.04.467325

**Authors:** Kathryn R. Taylor, Tara Barron, Helena Zhang, Alexa Hui, Griffin Hartmann, Lijun Ni, Humsa S. Venkatesh, Peter Du, Rebecca Mancusi, Belgin Yalçin, Isabelle Chau, Anitha Ponnuswami, Razina Aziz-Bose, Michelle Monje

**Affiliations:** Department of Neurology, Stanford University, Stanford, California USA; Department of Pediatrics, Stanford University, Stanford, California USA; Department of Pathology, Stanford University, Stanford, California USA; Department of Neurosurgery, Stanford University, Stanford, California USA

**Keywords:** Cancer Neuroscience, glioma, synapse, synaptic plasticity, AMPAR, BDNF, TrkB, DIPG, GBM

## Abstract

The nervous system plays an increasingly appreciated role in the regulation of cancer. In malignant gliomas, neuronal activity drives tumor progression not only through paracrine signaling factors such as neuroligin-3 and brain-derived neurotrophic factor (BDNF)^1–3^, but also through electrophysiologically functional neuron-to-glioma synapses^4–6^. Malignant synapses are mediated by calcium-permeable AMPA (α-amino-3-hydroxy-5-methyl-4-isoxazole propionic acid) receptors in both pediatric and adult high-grade gliomas^4, 5^, and consequent depolarization of the glioma cell membrane drives tumor proliferation^4^. The nervous system exhibits plasticity of both synaptic connectivity and synaptic strength, contributing to neural circuit form and functions. In health, one factor that promotes plasticity of synaptic connectivity^7, 8^ and strength^9–13^ is activity-regulated secretion of the neurotrophin BDNF. Here, we show that malignant synapses exhibit similar plasticity regulated by BDNF-TrkB (tropomyosin receptor kinase B) signaling. Signaling through the receptor TrkB^14^, BDNF promotes AMPA receptor trafficking to the glioma cell membrane, resulting in increased amplitude of glutamate-evoked currents in the malignant cells. This potentiation of malignant synaptic strength shares mechanistic features with the long-term potentiation (LTP)^15–23^ that is thought to contribute to memory and learning in the healthy brain^22 24–27 28, 29^. BDNF-TrkB signaling also regulates the number of neuron-to-glioma synapses. Abrogation of activity-regulated BDNF secretion from the brain microenvironment or loss of TrkB in human glioma cells exerts growth inhibitory effects *in vivo* and in neuron:glioma co-cultures that cannot be explained by classical growth factor signaling alone. Blocking TrkB genetically or pharmacologically abrogates these effects of BDNF on glioma synapses and substantially prolongs survival in xenograft models of pediatric glioblastoma and diffuse intrinsic pontine glioma (DIPG). Taken together, these findings indicate that BDNF-TrkB signaling promotes malignant synaptic plasticity and augments tumor progression.

---

Gliomas are the most common primary brain cancers in both children and adults^30^. Glioma progression is robustly regulated by interactions with neurons^1–5^, including tumor initiation^3^ and growth^1, 3^. Neuron-glioma interactions include both paracrine factor signaling^1, 3^ and electrochemical signaling through AMPAR-mediated neuron-to-glioma synapses^4, 5^. Synaptic integration of high-grade gliomas such as glioblastoma and DIPG into neural circuits is fundamental to cancer progression in preclinical model systems^4, 5^. We hypothesized that gliomas may recruit mechanisms of adaptive neuroplasticity to elaborate and reinforce these powerful neuron-glioma interactions.

## Strength of Glutamatergic Currents in Malignant Glioma

In neurons, activity-regulated ^24, 25, 31^ plasticity of synaptic strength dynamically modulates neural circuit function^32^ and these synaptic changes are thought to underlie learning and memory^28 29^. One form of long-term potentiation (LTP) of synaptic strength involves increased AMPAR channel insertion in the postsynaptic membrane^33, 34^. Glutamatergic neurotransmission through NMDAR (N-methyl-D-aspartate receptor) channels and consequent calcium signaling can increase AMPAR channel trafficking to the postsynaptic membrane^15–23^ (Figure 1a). Another activity-regulated mechanism that can promote LTP is BDNF-TrkB signaling, which similarly recruits calcium signaling pathways and promotes AMPAR channel trafficking to the post-synaptic membrane^9–13^. Hypothesizing that gliomas may exhibit plasticity of the recently discovered glutamatergic synapses^4–6^ formed between presynaptic neurons and post-synaptic glioma cells, we explored single cell transcriptomic datasets to elucidate potential mechanisms of synaptic plasticity. Pediatric gliomas express very low levels of NMDAR subunits^4^, but express high levels of TrkB (*NTRK2*) in malignant cells (Extended Data Fig. 1, Supplementary Note 1, Supplementary Table S1). Unlike adult glioblastoma^35^, pediatric high-grade gliomas, such as DIPG, do not express *BDNF* ligand (Extended Data Fig. 1). We therefore tested the hypothesis that BDNF-TrkB signaling could induce plasticity of the malignant synapse, focusing on pediatric high-grade gliomas to specifically probe interactions between the neural microenvironment and the tumor. We performed whole-cell patch clamp electrophysiology of glioma cells xenografted to the hippocampus in an acute slice preparation (Figure 1b, c). In response to transient and local glutamate application, voltage-clamp recordings of xenografted glioma cells demonstrated inward currents that increased in amplitude after BDNF incubation (Figure 1d, e, f). This effect of BDNF on glutamate-evoked inward currents illustrates the potential for plasticity of the malignant postsynaptic response.

**Fig. 1.**
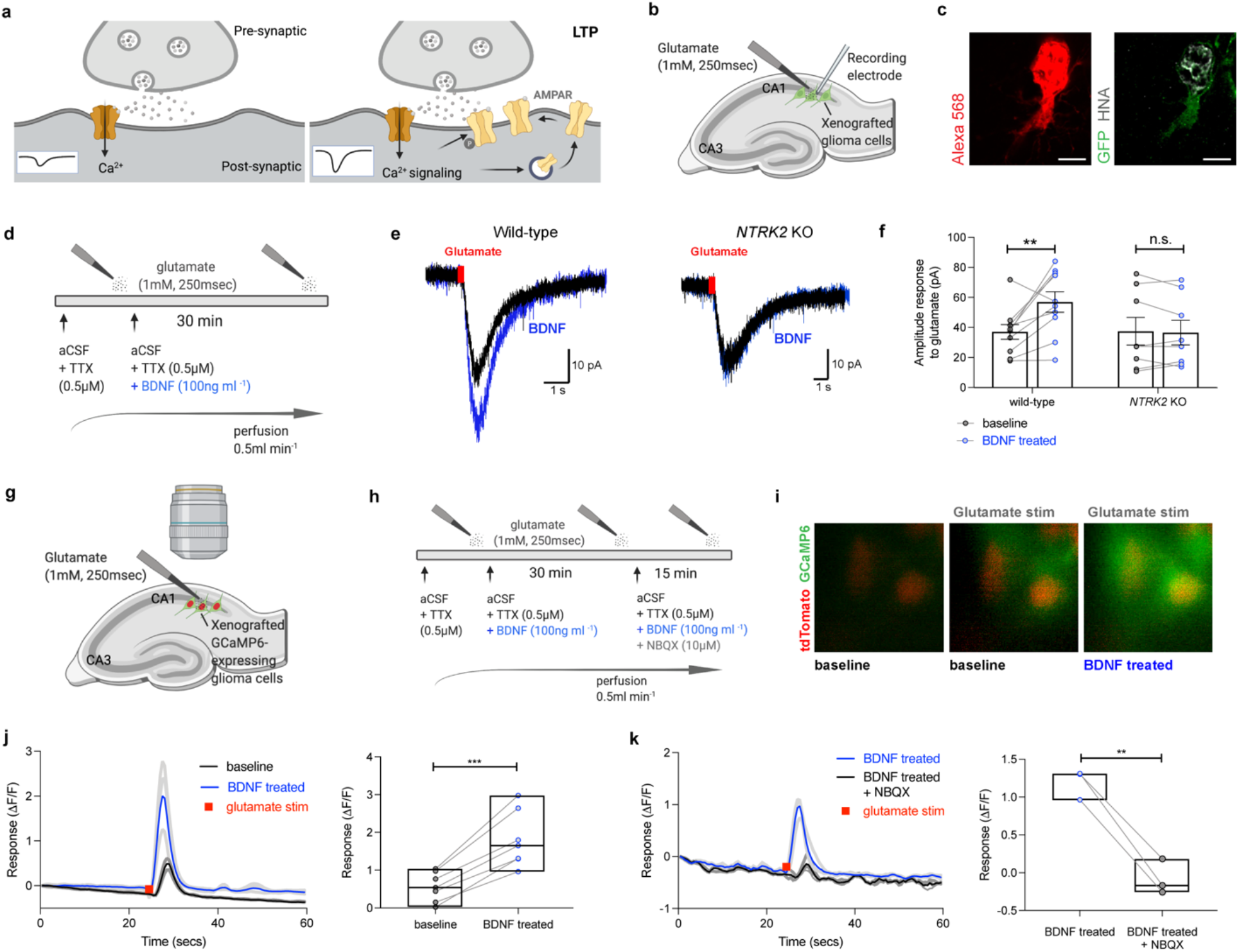
BDNF-TrkB signaling increases glutamatergic current amplitude in glioma. **a**, Simplified schematic outlining long-term potentiation via AMPAR channel trafficking to the postsynaptic density. NMDA-meditated LTP is the most extensively studied form of postsynaptic plasticity in CA1 hippocampal neurons. Activation of NMDAR channels following AMPAR-mediated cell depolarization allows calcium to enter the cell, inducing downstream activation of calcium pathway signaling^20^. Several kinases, such as CAMKII/PKC/MAPK have been demonstrated to alter AMPAR channel levels at the postsynaptic density^29^. **b**, Electrophysiological model of GFP+ glioma cells (green) xenografted in mouse hippocampal CA1 region to receive local and transient glutamate puff. **c**, Left, representative image of Alexa 568 (red)-filled GFP+ glioma cell post whole-cell patch clamp recording. Right, co-labelled with GFP (green) and HNA (grey). Scale bar = 5µm. **d**, Experimental outline of whole cell patch-clamp electrophysiological recordings in glioma cells. Hippocampal slices were perfused with ACSF containing 0.5 µM tetrodotoxin (TTX), and response to a local puff (250msec) application of 1 mM glutamate was recorded from xenografted glioma cells before and after perfusing with 100ng/ml BDNF for 30 min. **e**, Representative traces of glutamate-evoked inward currents in patient-derived glioma xenografted cells, Left wild-type glioma cells inward current response to glutamate puff, without (black) and with perfusion of aCSF containing 100ng/ml BDNF recombinant protein for 30mins (blue). Right, recordings of CRISPR-deleted *NTRK2* knockout glioma cells. **f**, Quantification of data in **e** (*n* = 10 wild-type cells, 6 mice and 8 knockout cells, 6 mice). **g**, Model of calcium imaging of tdTomato nuclear tag; GCaMP6s-expressing glioma cells xenografted into the mouse hippocampal region. **h**, Experimental outline of calcium transient recordings in patient-derived glioma cell xenografts. **i**, In-situ imaging of calcium transients in glioma cells evoked by local glutamate puff (1mM, 250msecs) at baseline (middle) and after perfusion with 100ng/ml BDNF in aCSF for 30mins. Green denotes glioma GCaMP6s fluorescence and red denotes tdTomato nuclear tag. **j**, Left, representative traces of glioma GCaMP6s intensity during local, transient glutamate application (red). GCaMP6s intensity response shown in three individual glioma cells (dark grey) to glutamate puff at baseline (average: black) and after BDNF perfusion (individual cells: light grey, average intensity: blue). Right, individual GCaMP6s cell response to glutamate puff at baseline and after BDNF application (*n* = 7 cells, 3 mice). **k**, Left, representative traces of glioma GCaMP6s intensity in the presence of BDNF. Response to glutamate application (red) were recorded with BDNF perfusion only (individual cells: light grey, average: blue) or with the addition of 10µm NBQX (individual cells: dark grey, average: black). Right, response of individual GCaMP6s cells to glutamate puff with BDNF application, in the presence and absence of NBQX. (*n*= 3 cells, 2 mice). Data are mean ± s.e.m. **P< 0.01, ***P<0.001, two-tailed paired Student’s *t*-test.

To confirm that glioma cell TrkB activation alters the glutamate-mediated currents, we used CRISPR technology to delete *NTRK2* from human glioma cells (referred to as *NTRK2* knockout (KO)). The knockout was a direct deletion within exon 1 of *NTRK2* and produced a ∼ 80% decrease in TrkB protein levels (Extended Data Fig. 2). This reduction of TrkB expression in the glioma cells blocked the BDNF-induced increase in glutamate-evoked inward current compared to *NTRK2* WT, Cas9-control glioma xenografts (Figure 1e, f), demonstrating the BDNF-TrkB signaling pathway modulates the strength of the glutamate-mediated currents in the tumor cells.

To explore the intracellular consequences of BDNF-induced current amplification we performed in situ calcium imaging of xenografted glioma cells that express the genetically encoded calcium indicator GCaMP6s (Figure 1g). As expected^4^, local glutamate application induced calcium transients in glioma cells (Figure 1h, i). The intensity of the glutamate-induced calcium transient was increased by BDNF incubation (Figure 1i, j). Glioma calcium transients evoked by glutamate were blocked by the AMPAR channel inhibitor, NBQX (Figure 1k), consistent with the known role for AMPAR channels mediating glutamatergic signaling in glioma^4, 5^. Taken together, these data illustrate that BDNF increases the strength of the glutamate-evoked response.

## Trafficking of AMPA Receptors in Malignant Glioma

In healthy neurons, BDNF activation of TrkB increases the levels and trafficking of AMPAR channels to the postsynaptic membrane via the PKC/CAMKII calcium signaling pathway (Figure 2a)^10, 36^. Given the findings above that BDNF increased AMPAR-mediated inward currents and calcium transients in glioma, we next investigated the effect of BDNF on AMPAR channel trafficking to the cell membrane. Glioma cell surface proteins were captured using biotinylation with avidin pull-down and probed for levels of AMPAR subunits. Consistent with the hypothesis that BDNF increases AMPAR channel trafficking to the glioma cell membrane, BDNF treatment increased glioma cell surface expression of the AMPAR subunit GluA4, compared to vehicle-treated control glioma cells (Figure 2b).

**Fig. 2.**
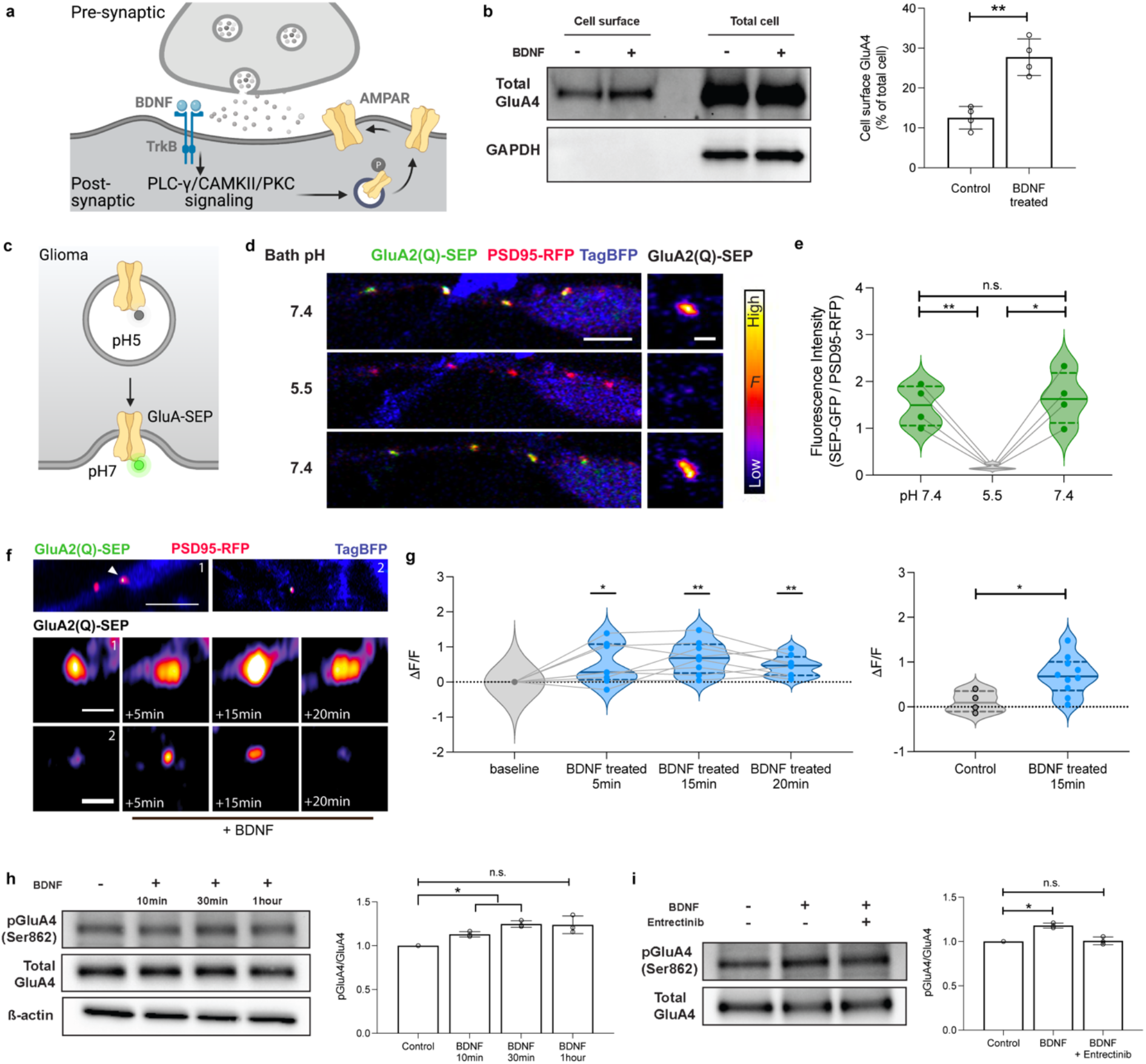
BDNF regulates trafficking of AMPAR to the glioma postsynaptic membrane. **a**, Schematic depicting AMPAR channel trafficking downstream of BDNF-TrkB signaling. BDNF-TrkB induces activation of Ca^2+^/calmodulin-dependent kinase II (CAMKII) and protein kinase C (PKC), which phosphorylates AMPAR subunits and increases synaptic delivery^36^. **b**, Western blot analysis of cell membrane surface and total cell protein from SU-DIPGVI cells treated with 100nM BDNF for 30mins. Surface proteins were labelled with Biotin and extracted from total protein with Avidin bead pull-down. Right, quantification of % of biotinylated cell surface proteins from total (*n* = 4 biological replicates). **c,** Super ecliptic pHluorin tagged GluA subunit (GluA-SEP). **d-e,** Validation of pHluorin. **d,** Left, confocal image of a glioma cell expressing the calcium permeable AMPAR subunit tagged with SEP (GluA2(Q)-SEP), PSD95-RFP and whole cell TagBFP, in co-culture with neurons. Right, individual representative puncta of GluA2(Q)-SEP. Scale bars = 5µm and 1µm, respectively. Cells were perfused with pH 7.4 aCSF followed by exposure to aCSF containing membrane impermeable acid at pH 5.5 and then returned to pH 7.4. **e,** Quantification of GluA2(Q)-SEP puncta intensities before, during and after acidic aCSF exposure, as a ratio to PSD95-RFP puncta intensity (*n* = 4 puncta, one cell). **f**, confocal images of two representative GluA2(Q)-SEP; PSD95-RFP, TAG-BFP expressing glioma cells in co-culture with neurons. Co-localized puncta intensity was measured over a time course with BDNF (100nM) perfusion. **g,** Left, quantification of GluA2(Q)-SEP / PSD95-RFP puncta intensities over time with BDNF as compared to initial baseline intensity (*n* = 8 puncta, 6 cells). Right, quantification of one 15-minute time point with control cells (vehicle, *n* = 4 puncta, 2 cells) vs BDNF treatment (100nM, *n* = 8 puncta, 6 cells). **h,** Representative Western blot analysis of primary patient-derived glioma culture, SU-DIPGVI, treated with 100nM recombinant BDNF at several timepoints using indicated antibodies. Right, quantification of the phospho-immunoblots ratio to corresponding total protein levels and normalized to vehicle treated control (*y* axis is in arbitrary units, *n* = 3 biological replicates). **i**, Representative Western blot analysis of 100nM BDNF treated glioma cells (as in **h**,) at 30mins with and without entrectinib (5µM). Right, quantification of phospho-immunoblots (as in **h**, *y* axis is in arbitrary units, *n* = 3 biological replicates). Data are mean ± s.e.m. *P< 0.05, **P< 0.01, two-tailed paired Student’s *t*-test for **b**, **e**, One sample t and Wilcoxon test for g (left panel), **h** and **i**. Unpaired t-test for **g** (right panel).

To further test this hypothesis, we leveraged pHluorin technology for live imaging of AMPAR trafficking within glioma cells co-cultured with neurons. We expressed GluA2 AMPAR subunit tagged to a pH sensitive GFP (pHluorin) in glioma cells and then performed high-resolution confocal live imaging of these AMPAR channels within glioma cultures^37^. We additionally expressed PSD95 tagged to RFP in glioma cells in order to confirm the localization of the AMPAR channels to the glioma postsynaptic site. Super ecliptic pHluorins (SEPs) fluoresce when the N-terminus of the channel moves from the acidic pH within the trafficking vesicle, to the neutral pH on the outside of the cell membrane (Figure 2c). To validate the pHluorin strategy, we confirmed that exposure of GluA2(Q)-SEP:PSD95-RFP-expressing glioma cells to acidic medium (pH 5.5) quenched the signal, demonstrating that the majority of the AMPAR fluorescent signal at the postsynaptic puncta is from membrane-bound GluA2 (Figure 2d, e). Time course imaging of individual puncta demonstrated that BDNF exposure elicits an increase in the postsynaptic levels of AMPAR channels on the glioma cells (Figure 2f, g). Taken together, these findings indicate that BDNF-TrkB signaling increases trafficking of AMPAR channels to the cell membrane, accounting for the increased glutamate-evoked currents described above.

We next explored the signaling mechanisms of BDNF in patient-derived pediatric glioma cells. Several signaling pathways are known to be activated upon BDNF binding to the TrkB receptor, and the expression of different TrkB splice variants can alter the function of the receptor^38^. Using the publicly available transcript data of pediatric glioma, we compared the expression of the *NTRK2* splice variants and found that whilst gliomas do express full-length TrkB, the tumors have a higher expression of truncated TrkB, as has been previously described^39^ (Extended Data Fig. 3). Western blot analysis of glioma cells demonstrates that BDNF recombinant protein treatment activates three main signaling cascades; MAPK, PI3K and calcium signaling (Extended Data Fig. 4). In neurons, BDNF increases AMPAR trafficking via CAMKII/PKC calcium signaling^10^, kinases known to play roles in neuronal LTP^19, 22, 40^. MAPK and PI3K have also been shown to play a role in AMPAR transmission^41^. Signaling-induced posttranslational modifications, such as phosphorylation, instruct the activation, conductance and trafficking of AMPAR subunits^42^. We found that BDNF exposure increases phosphorylation of the subunit GluA4 at ser862, a site shown to facilitate the delivery of the channel to the postsynaptic density in glioma cells (Figure 2h)^43^. Treatment with a pan-Trk inhibitor, Entrectinib, abrogated this BDNF-induced increase in glioma GluA4 phosphorylation (Figure 2i). In contrast to these protein phosphorylation and trafficking effects of BDNF, pediatric glioma cells exhibited few gene expression changes in response to BDNF exposure, with the exception of VGF (Extended Data Fig. 5) -a gene recently shown to be regulated by BDNF in adult GBM as well^35^.

## Plasticity of Synaptic Connectivity

A subset of glioma cells engage in synapses^4, 5^ and accordingly we observed a subset xenografted glioma cells that exhibit an inward current in response to glutamate using patch clamp electrophysiology as above. *NTRK2* KO tumors exhibited a reduced number of cells that responded to glutamate application with an inward current (Figure 3a, b). We hypothesized that *NTRK2* loss in glioma cells may alter the level of malignant synaptic connectivity.

**Fig. 3.**
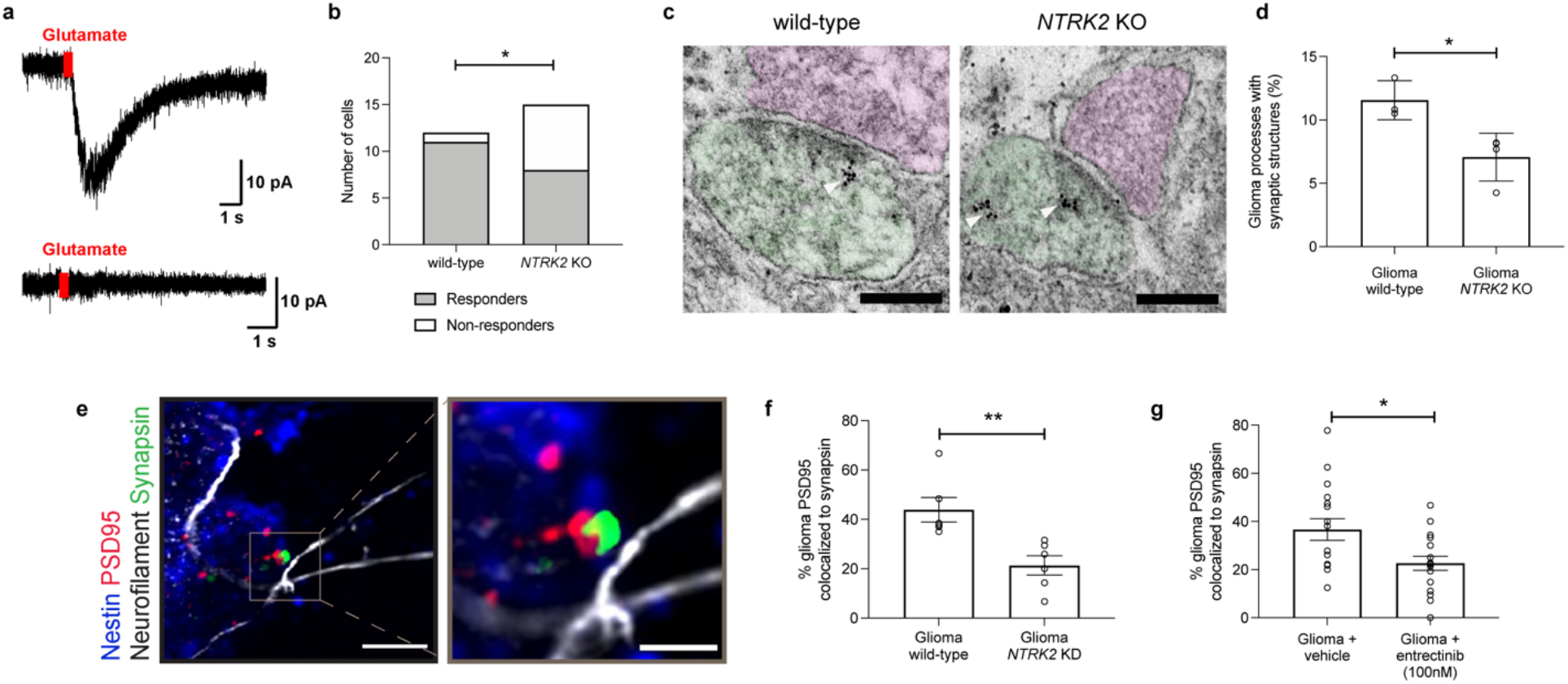
TrkB regulates neuron-glioma connectivity. **a,** Representative electrophysiological traces of glutamate-evoked inward currents in patient-derived glioma xenografted cells (as in Figure 1b), with cells that respond to glutamate puff (top) and those that have no current change (bottom). **b,** Quantification of data in **a** (*n* = 12 wild-type cells, 6 mice and 15 knockout cells, 7 mice). **c,** Immuno-electron microscopy of patient-derived DIPG cells SU-DIPG-VI wild-type (left) and *NTRK2* KO (right) xenografted into the mouse hippocampus. Arrowheads denote immuno-gold particle labelling of GFP. Synapses confirmed as GFP+ postsynaptic density in glioma cells (colored green), a synaptic cleft and clustered synaptic vesicles in opposing presynaptic neuron (colored magenta). Scale bar = 2µm. **d,** Quantification of identified synapses in **c** for mice harboring wild-type and *NTRK2* KO tumors (*n* = 3 mice/group). **e**, Confocal images of neurons co-cultured with PSD95-RFP labelled wild-type and *NTRK2* KO glioma cells. White denotes neurofilament (axon); green denotes nestin staining (glioma cell process); blue denotes synapsin (presynaptic puncta). Scale bar = 5µm. **f**, Quantification of the colocalization of postsynaptic glioma-derived PSD95-RFP with neuronal presynaptic synapsin in co-cultures of wild-type (*n* = 6 cells, 3 coverslips from 3 biological replicates), or *NTRK2* KO glioma cells (SU-DIPG-VI, *n* = 6 cells, 3 coverslips from 3 biological replicates). **g,** Quantification of the colocalization of postsynaptic glioma-derived PSD95-RFP with neuronal presynaptic synapsin in neuron co-cultures with SU-DIPG-VI glioma cells treated with vehicle or entrectinib (100nM) (*n* = 18, cells, 6 coverslips from 3 biological replicates). Data are mean ± s.e.m. **P< 0.01, ***P<0.001, * P< 0.05, Fisher’s exact test for **b**. Two-tailed unpaired Student’s t-test for **d**, **f** and **g**.

To investigate whether BDNF-TrkB signaling regulates the number of neuron-glioma synaptic connections we performed immuno-electron microscopy in wild-type and *NTRK2* KO patient-derived glioma xenografts expressing green fluorescent protein (GFP). Using immuno-electron microscopy with immunogold labeling of GFP+ cells to unambiguously identify the malignant cells, we identified fewer neuron-to-glioma synaptic structures in the *NTRK2* KO tumors compared to wild-type tumors (Figure 3c, d). Glioma cultures expressing RFP-tagged PSD95 were subjected to shRNA knockdown of *NTRK2* (KD) and cultured in the presence of neurons (Extended Data Fig. 2). We found that co-culture of *NTRK2* KD glioma cells exhibited fewer structural synapses with neurons evident as co-localized neuronal presynaptic puncta (synapsin) with glioma postsynaptic puncta (PSD95-RFP) in comparison to *NTRK2* wild-type glioma cells (Figure 3e, f). A similar reduction in structural neuron-to-glioma synapses was seen upon the addition of the pan-Trk inhibitor, entrectinib, to neuron-glioma co-cultures (Figure 3g). Taken together, these data demonstrate that BDNF-TrkB signaling modulates neuron-glioma synaptic connectivity.

## BDNF regulates glioma proliferation in the context of neurons

BDNF alone can increase glioma proliferation^1, 3, 35^, albeit not as robustly as other neuron-glioma signaling mechanisms^1^. In contrast to adult glioblastoma^35^, single cell transcriptomic studies demonstrate that pediatric glioma cells do not express BDNF ligand (Extended Data Figure 1) and thus the chief source of BDNF is the brain microenvironment. Testing the effects of BDNF alone on glioma proliferation *in vitro*, we found that the addition of recombinant BDNF (100 nM) increases pediatric glioma (DIPG) cell proliferation from a rate of ∼20% to ∼30%. This effect is completely abrogated -as expected - with CRISPR knockout of *NTRK2* (Figure 4a,b). Similar results of a small increase in proliferation are observed in a range of patient-derived glioma cultures exposed to BDNF, including thalamic DMG and pediatric cortical glioblastoma (Extended Data Fig. 6).

**Fig. 4.**
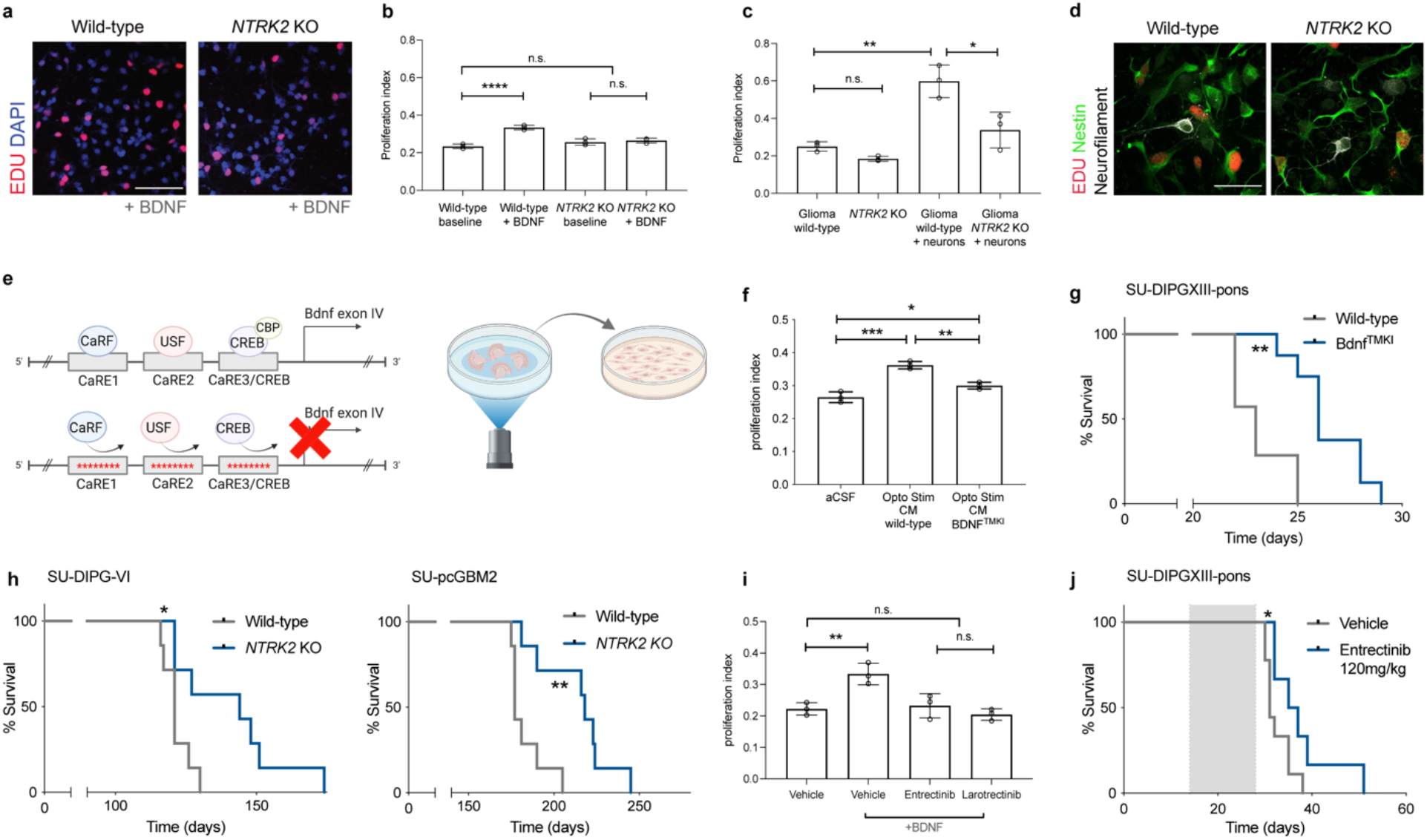
Activity-regulated BDNF promotes glioma progression. **a,** Confocal image of wild-type and *NTRK2* KO patient-derived glioma cultures (SU-DIPG-VI) cultured with and without 100µM of BDNF protein. Red denotes EdU incorporation and DAPI marked nuclei in blue. Scale bar =100µm **b,** Proliferation index of cultures in **a** with and without BDNF treatment. **c,** Proliferation index of wild-type and *NTRK2* KO glioma cultures (SU-DIPG-VI) cultured alone or with neurons. **d,** Representative image of wild-type and *NTRK2* KO glioma cells (SU-DIPG-VI) co-cultured with neurons quantified in **c**. Green denotes nestin positive glioma cells; white denotes Neurofilament (neurons); red denotes EdU (proliferative marker). Scale bar = 50µm. **e,** Left, schematic of BDNF^TMKI^ mouse, which lacks activity-regulated *BDNF* expression. Right, collection of conditioned medium (CM) from optogenetically stimulated acute cortical slices. **f,** Proliferation index of SU-DIPG-VI cells exposed to wild-type or BDNF^TMKI^ CM. **g,** Kaplan-Meier survival curves of wild-type and BDNF^TMKI^ mice bearing patient-derived orthotopic xenografts of SU-DIPGXIII-P* (*n* = 7 wild-type mice, 8 BDNF^TMKI^ mice). **h,** Survival curves of mice bearing wild-type and *NTRK2* KO orthotopic xenografts for two patient-derived pediatric glioma models (SU-pcGBM2, H3WT pediatric cortical glioblastoma model and SU-DIPG-VI, DIPG model; *n* = 7 mice in each group). **i,** Proliferation index of SU-DIPG-VI cells treated with 100µM of BDNF protein in the presence of pan-Trk inhibitors, entrectinib and larotrectinib at 500nM (*n* = 3 biological replicates per group for **b, c, f, i**). **j,** Survival curves of pontine-injected SU-DIPGXIII-P* patient-derived orthotopically xenografted mice treated with 120mg/kg entrectinib vs vehicle treated controls. Data are mean ± s.e.m. *P< 0.05, **P< 0.01, ***P<0.001, one way ANOVA with Tukey’s post hoc analysis for **b**, **c, f** and **i**. Two-tailed log rank analysis for **g**, **h** and **j**.

In contrast to the relatively weak mitogenic effect of BDNF on glioma cells alone, co-culture with neurons elicits a robust increase in proliferation rate from ∼20% to ∼60% EdU+ glioma cells, underscoring the powerful effects of neurons on glioma proliferation that includes neuroligin-3 signaling and neuron-to-glioma synaptic mechanisms^1, 3–5^. We sought to investigate the relative contribution of BDNF-TrkB signaling in neuron-glioma interactions using neuronal co-culture with WT or *NTRK2* KO glioma cells. A reduction in pediatric glioma proliferation from *NTRK2* loss was not observed in the absence of neurons (Figure 4c, d), consistent with the lack of BDNF ligand expression in pediatric glioma cells (Extended Data Fig. 1). However, TrkB loss in glioma cells co-cultured with neurons resulted in a stark reduction in neuron-induced proliferation, decreasing the proliferation rate from ∼60% to ∼30% EdU+ glioma cells. This reduction was disproportionate to the loss accounted for by BDNF mitogenic signaling alone (Figure 4b). The magnitude of the change in glioma proliferation elicited by *NTRK2* loss in response to BDNF recombinant protein compared to that elicited by co-culture with neurons (Figure 4b, d, 10.02% ± 1.2 vs 26.06% ± 2.3 *P* = 0.0004) suggests that BDNF is playing a more complex role in neuron-glioma interactions than simply acting as an activity-regulated growth factor. This is consistent with the idea that BDNF-TrkB signaling regulates neuron-to-glioma synaptic strength and connectivity, as demonstrated above.

## Activity-dependent BDNF promotes tumor progression

We next tested the necessity of neuronal activity-regulated BDNF for glioma growth using a genetically engineered mouse model deficient in activity-induced expression of Bdnf (*Bdnf^TMKI^*; TMKI, triple-site mutant knockin^44^). This mouse model expresses baseline levels of BDNF, but does not exhibit activity-regulated increase in BDNF expression and secretion due to loss of the CREB binding site in the *Bdnf* promotor^44^. First, to confirm the relative contribution of activity-regulated BDNF ligand to the mitogenic effect of activity-regulated secreted factors, we crossed the *Bdnf^TMKI^* mouse to mice expressing the excitatory opsin channelrhodopsin in deep cortical projection neurons (Thy1::ChR2, Figure 4e), then optogenetically stimulated cortical slices from Thy1::ChR2-expressing mice with *Bdnf^TMKI^* or wild-type *Bdnf*. The effects of conditioned media (CM) from optogenetically-stimulated cortical slices on glioma cell proliferation was assessed using our well-validated experimental paradigm^1^. Exposure of glioma patient-derived cultures to CM from WT optogenetically stimulated slices increased proliferation rate, as we have previously shown^1^ (Figure 4f). Conditioned media collected from optogenetically stimulated *Bdnf^TMKI^* slices elicited a mildly reduced proliferative response compared to WT, indicating a small role for BDNF ligand in paracrine neuron-glioma interactions (Figure 4f), as expected^1^. Given the results in neuron-glioma co-culture described above, we would anticipate that loss of activity-regulated BDNF would hinder glioma growth *in vivo* to a greater extent. Concordantly, survival analysis of orthotopic pediatric glioma xenografts in WT and *Bdnf^TMKI^* mice demonstrated markedly increased survival in *Bdnf^TMKI^* mice lacking activity-regulated BDNF secretion (Figure 4g, Extended Data Fig. 7), concordant with the hypothesis that activity-regulated BDNF signaling influences glioma progression in the context of the brain microenvironment in important ways.

## Therapeutic targeting of TrkB in pediatric glioma

We next tested the effects of genetic or pharmacological blockade of TrkB on pediatric glioma growth. Mice were xenografted orthotopically with patient-derived cells in which *NTRK2* was wild type (Cas9 control) or had been CRISPR-deleted. Mice bearing orthotopic xenografts of *NTRK2* KO DIPG or *NTRK2* KO pediatric cortical glioblastoma exhibited a marked increase in overall survival compared with littermate controls xenografted with *NTRK2* WT cells (Figure 4h, Extended Data Fig. 7).

We next performed preclinical efficacy studies of pan-Trk inhibitors. Trk inhibitors have recently been developed for treatment of NTRK-fusion malignancies, including *NTRK* fusion infant gliomas^45–47^. Here, we tested the preclinical efficacy of these inhibitors in *NTRK* non-fusion gliomas like DIPG. We first confirmed that both entrectinib and larotrectinib abrogated glioma proliferation in response to BDNF ligand stimulation (Figure 4i). We then tested the ability of entrectinib to cross the blood brain barrier and demonstrated that systemic entrectinib (120 mg/kg PO) reduced TrkB phosphorylation and downstream ERK phosphorylation in brain tissue (Extended Data Fig. 8). Treatment of an aggressive patient-derived pediatric glioma (DIPG) orthotopic xenograft model with entrectinib increased overall survival compared to vehicle-treated controls (Figure 4j, Extended Data Fig. 7).

## DISCUSSION

Neurons form synapses with glioma cells via calcium permeable AMPA receptors^4, 5^ and the consequent membrane depolarization promotes glioma growth through voltage-sensitive mechanisms that remain to be fully elucidated^4^. Tumor cells form a network with each other through long processes called tumor microtubes connected by gap junctions^48–50^, and neuron-glioma electrochemical communication propagates through the glioma network through this gap junctional coupling^4, 5^ such that a single neuron-glioma synapse may affect numerous glioma cells. Here, we find that malignant synapses exhibit plasticity of both strength and number. Increased AMPAR channel trafficking to the glioma cell membrane mediates this plasticity of synaptic strength, recapitulating a mechanism of long-term potentiation (LTP) operational in healthy neurons that contributes to learning and memory^22, 24–29^. Whether other mechanisms of synaptic plasticity^32^ occur in glioma remain to be determined in future work.

Neuronal activity promotes glioma progression through paracrine^1–3^ and synaptic^4, 5^ signaling mechanisms. The findings here illustrate that neuronal activity-regulated factors such as BDNF not only directly promote glioma growth^1, 3, 35, 39^, but can also further reinforce neuron-glioma interactions. This potential for plasticity of malignant synaptic strength and connectivity raises a number of questions about neuron-to-glioma network evolution over the disease course. Could this activity-dependent reinforcement of neuron-glioma interactions and increased synaptic integration contribute to treatment resistance later in disease course? Could experience and activity patterns in certain patients contribute to neuroanatomical location of disease progression? Limiting malignant network elaboration by targeting malignant synaptogenesis and plasticity may be crucial for disease control, a concept supported by the therapeutic potential of disrupting BDNF-TrkB signaling demonstrated here.

Gliomas hijack processes of neural plasticity and integrate into neural networks in complex and dynamic ways, leveraging mechanisms that normally regulate neural circuit establishment during development and plasticity contributing to cognition in the healthy brain. Understanding and targeting neural circuit mechanisms in glioma will be critical for effective treatment of these deadly brain cancers.

## Supplementary Figures

**Extended Data Fig. 1.**
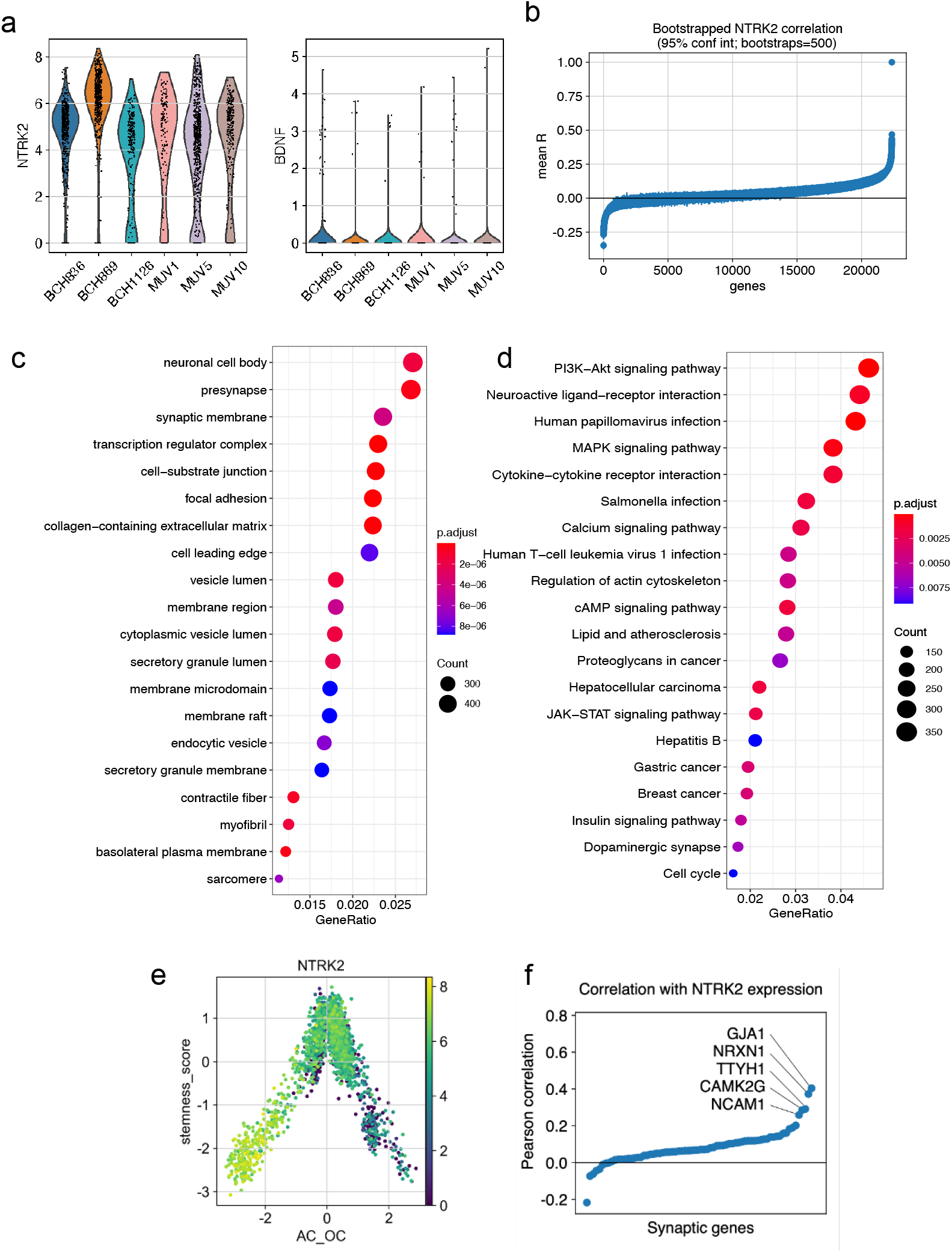
NTRK2 and BDNF expression in pediatric gliomas. **See Supplemental Note 1. a**, *NTRK2* (left) and *BDNF* (right) expression in H3K27M^+^ DMG malignant single cells from primary human biopsy single-cell transcriptomic data (scRNASeq, *n* = 2,259 cells, 6 study participants). (*P* <0.0001 for *NTRK2* vs *BDNF* expression in each tumor sample, value estimated using paired t-test). **b**, Plot of expression correlation analysis, performed using the bootstrap method on the H3K27M^+^ DMG scRNASeq data, to identify genes which correlate with *NTRK2* expression. **c,** Gene Ontology (GO) cellular compartments enriched for the top 92 genes correlated with *NTRK2* expression (identified in **b**). **d,** KEGG analysis demonstrated key signaling pathways enriched for the top 92 genes correlated with *NTRK2* expression (identified in **b**). **e**, *NTRK2* expression level in malignant H3K27M^+^ malignant single cells plotted using the lineage (*x* axis) and stemness (undifferentiated to differentiated; *y* axis) scores. **f**, Pairwise Pearson correlation analysis of 73 synaptic-associated genes with expression of *NTRK2*, the genes with a Pearson correlation > 0.25 are highlighted.

**Extended Data Fig. 2.**
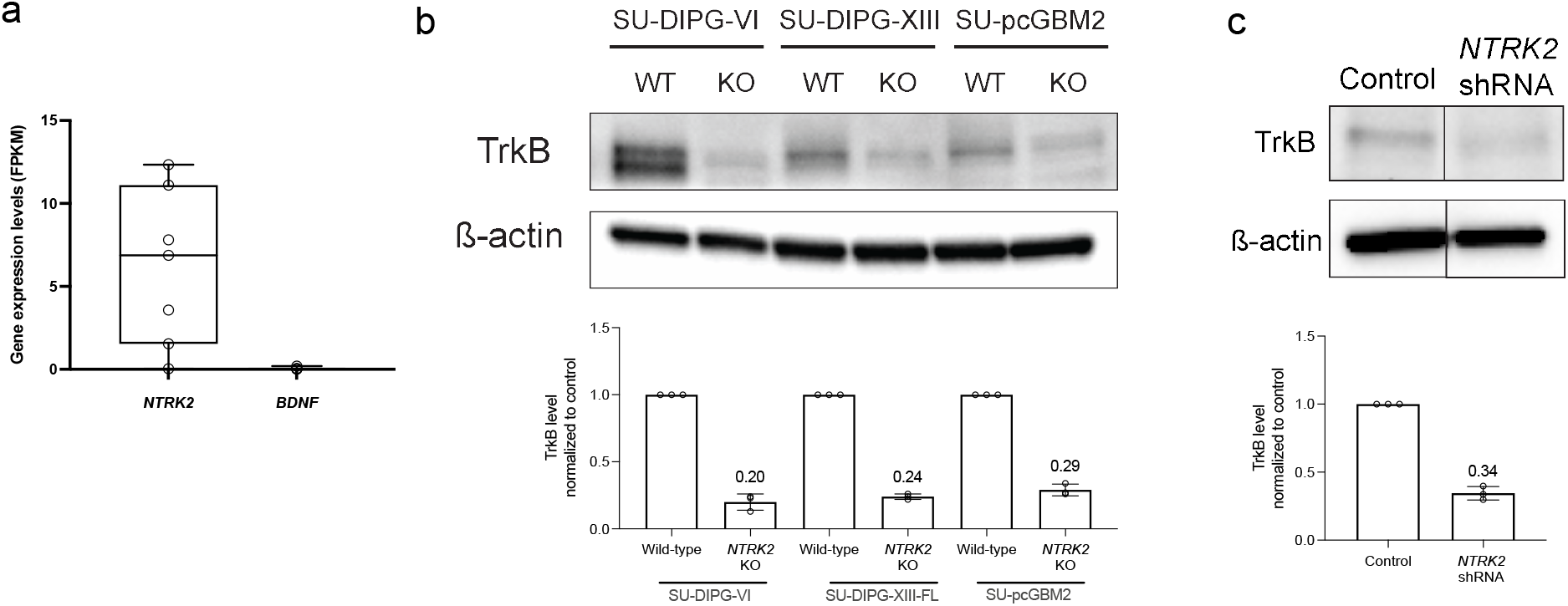
Genetic deletion of *NTRK2* from pediatric glioma patient-derived cultures. **a,** Analysis of previously published bulk RNA sequencing data of patient derived cultures (including SU-DIPG-VI, SU-DIPGXIII used in our experiments), confirmed the expression of *NTRK2* in these cultures (in Ext Data Fig. 1a), with little expression of BDNF (mean 6.180 vs 0.052 FPKM respectively, *P* value estimated by Wilcoxon matched pairs signed rank test). **b**, Top, representative western blot analysis of TrkB protein levels in wild-type, Cas9-control and *NTRK2* KO cultures (SU-DIPG-VI, SU-pcGBM2, SU-DIPGXIII-FL), using indicated antibodies. Bottom, quantification of western blot analysis with levels of TrkB normalized to total protein loading using ß-actin levels and compared to wild-type, Cas9-scramble control, cultures (*y* axis is in arbitrary units, *n* = 3 biological replicates). **c,** Top, representative western blot analysis of TrkB protein levels in control scramble shRNA and *NTRK2* shRNA knockdown cultures (SU-DIPG-VI), using indicated antibodies and analyzed as in **b,**. The samples were run together in the same western blot experiments, but had one additional sample in between from another shRNA knockdown culture that was not used in these experiments. Bottom, quantification of western blot analysis with levels of TrkB normalized to total protein loading using ß-actin levels and compared to wild-type, shRNA-scramble control, cultures (*y* axis is in arbitrary units, *n* = 3 biological replicates). Data are mean ± s.e.m. **P< 0.01, ***P<0.001, One sample *t*-test.

**Extended Data Fig. 3.**
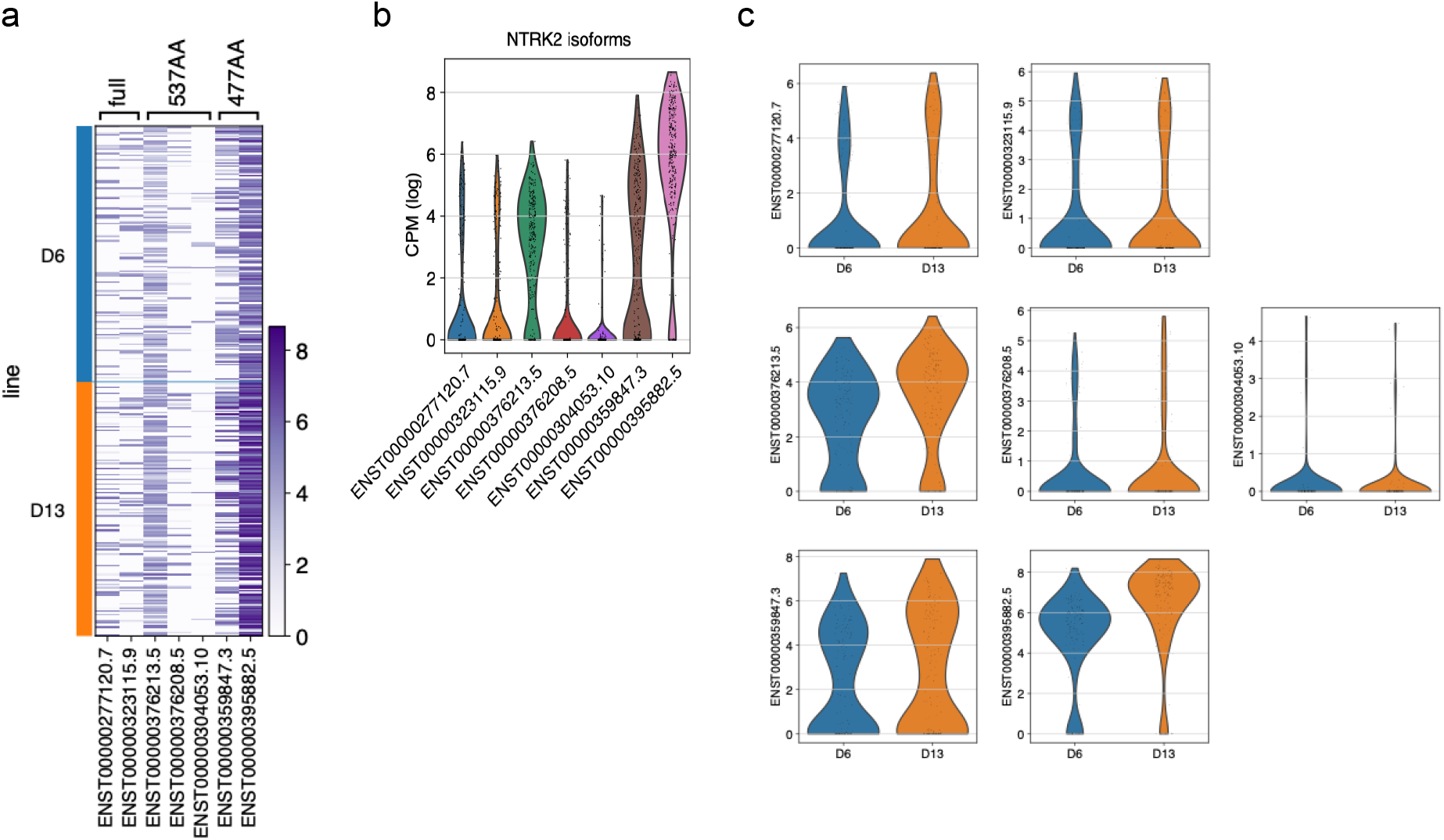
TrkB isoform expression in DIPG. **a,** Heatmap of single-cell RNASeq data of patient derived glioma xenograft models (SU-DIPG-VI and SU-DIPGXIII) cells (*n* = 321 cells, 4 mice) demonstrating relative expression of TrkB (*NTRK2*) isoforms; Full (ENST-277120.7 and ENST-323115.9) and Truncated (527aa ENST-376208.5 and ENST-376208.5; 477aa ENST-359847 and ENST-395882.5), with representative Ensembl codes depicted below. **b,** Violin plots of relative expression level of TrkB isoforms (depicted in **a,**) for both SU-DIPG-VI and SU-DIPGXIII cells combined, shown as log-transformed counts per million (CPM). **c,** Violin plots of relative expression of TrkB isoforms (depicted in **a,**) separated out by patient-derived xenograft model type (SU-DIPG-VI (D6) and SU-DIPGXIII (D13)). X-axis is in log-transformed counts per million (CPM).

**Extended Data Fig. 4.**
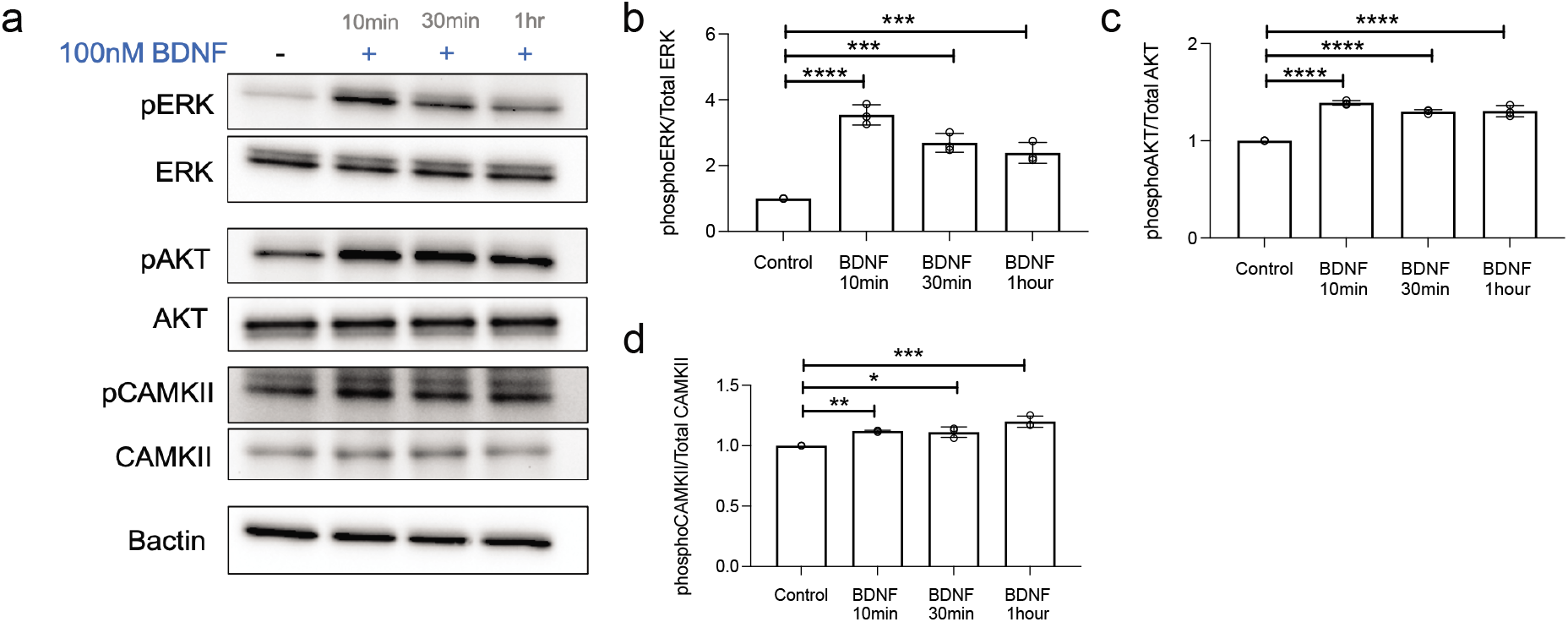
BDNF treatment induces PI3K, MAPK and CAMKII activation. **a,** Western blot of proteins from SU-DIPG-VI cells treated with BDNF recombinant protein (100nM) over a time course and probed for the indicated antibodies to demonstrate activation of downstream signaling pathways in comparison to untreated cells (vehicle only). **b,** Quantification of MAPK pathway activation in **a**, by comparing the ratio of the normalized phospho-ERK (T202/Y204) levels to corresponding total protein levels for BDNF treated cultures compared to control (*y*-axis is in arbitrary units, *n* = 3 biological replicates). **c,** Quantification of PI3K pathway activation in **a**, by comparing the ratio of the normalized phospho-AKT (S473) to corresponding total protein levels for BDNF treated cultures compared to control (*y*-axis is in arbitrary units, *n* = 3 biological replicates). **d,** Quantification of calcium pathway activation in **a**, by comparing the ratio of the normalized phospho-CAMKII (T286) to corresponding total protein levels for BDNF treated compared to control (*y* axis is in arbitrary units, *n* = 3 biological replicates). Data are mean ± s.e.m. *P< 0.05, **P< 0.01, ***P<0.001, one-way analysis of variance (ANOVA) with Tukey’s post hoc analysis.

**Extended Data Fig. 5.**
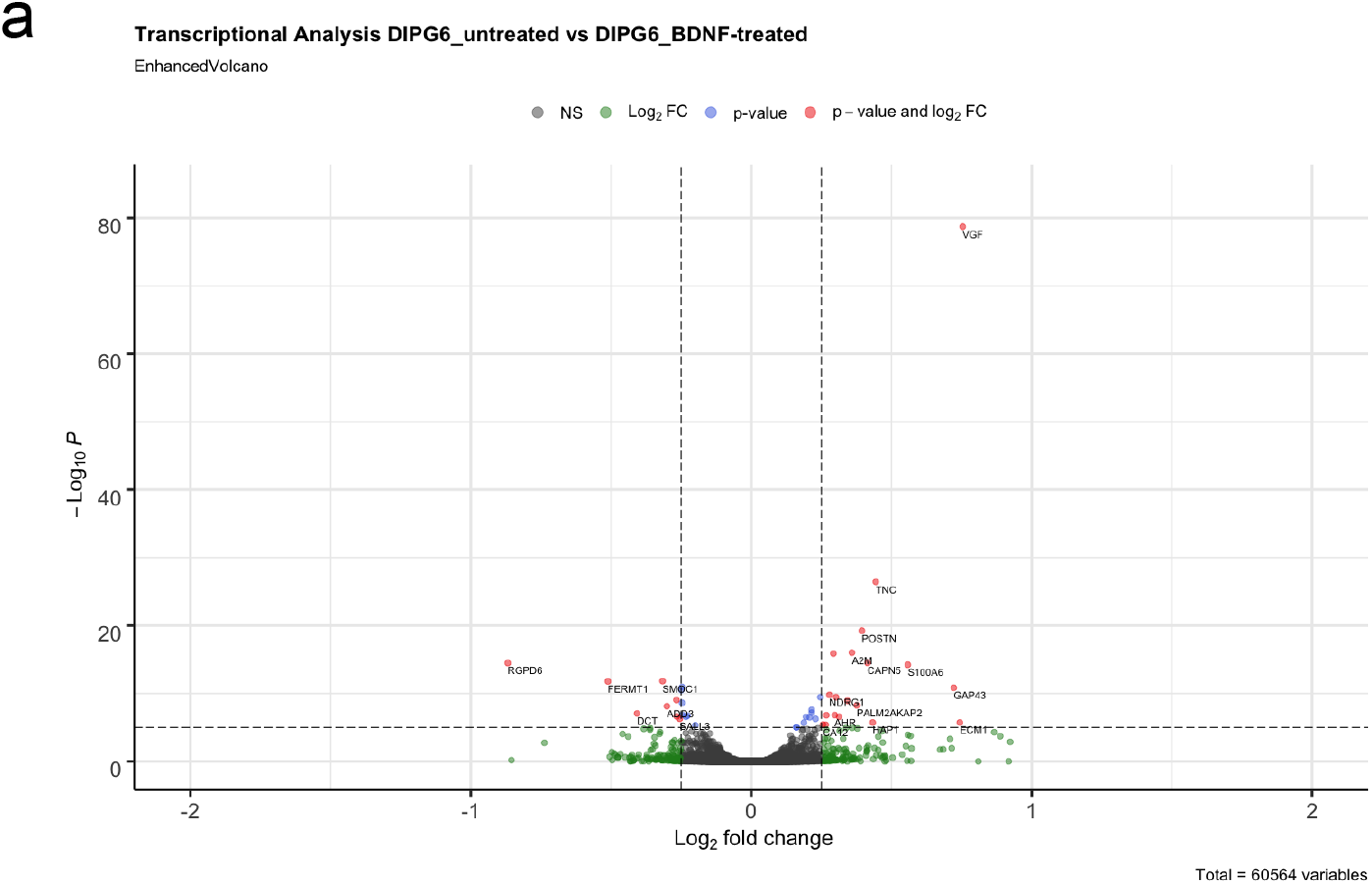
Gene expression changes induced by BDNF treatment in glioma. **a,** Scatterplot demonstrating gene expression changes in SU-DIPG-VI after 16 hours of treatment with and without BDNF recombinant protein (100nM) compared to control cells (vehicle treated). The x axis demonstrates log_2_ (fold change of BDNF compared to vehicle) and the x axis demonstrates Log_10_P of the gene expression level. Points shown in red represent genes showing statistically significant change (adjusted *P*<0.1, Benjamin-Hochnerg for multiple comparison testing).

**Extended Data Fig. 6.**
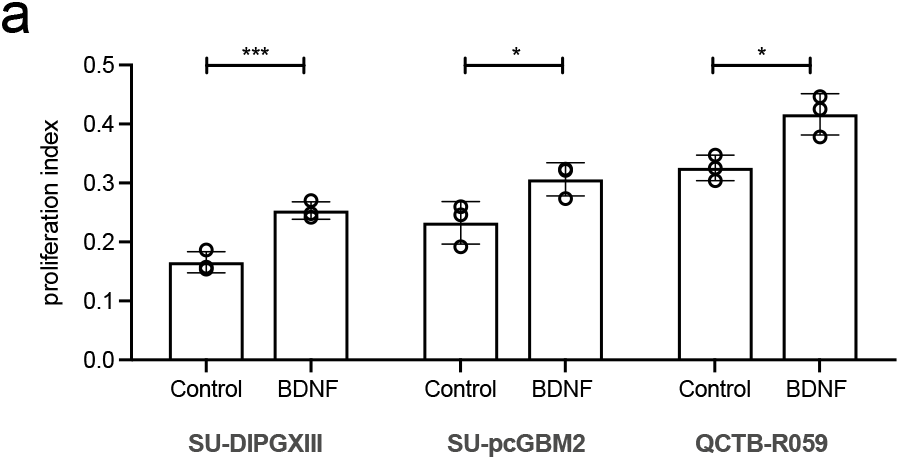
BDNF-induced proliferation in pediatric glioma. **a,** Proliferation index of DIPG (XIII-FL), cortical (pcGBM2) and thalamic (QCTB-R059) pediatric glioblastoma cultures treated with BDNF recombinant protein (100nM) compared to control cells (vehicle treated). Confocal images of the treated glioma cultures were taken (as in Figure 4a), with EdU+ cells counted as a proportion of total cells marked by DAPI nuclei staining (*n* = 3 biological replicates for all).

**Extended Data Fig. 7.**
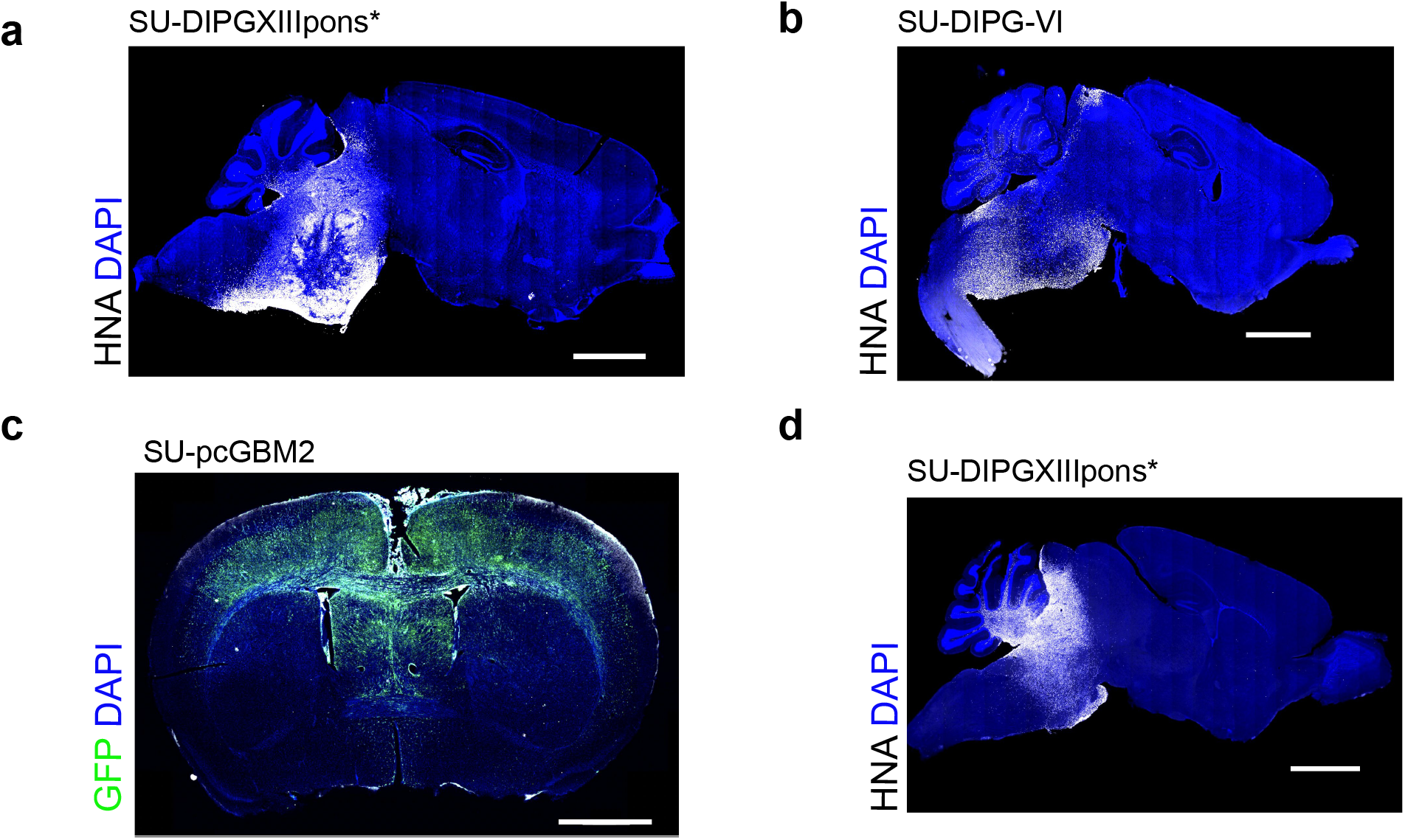
Orthotopic xenograft models used in survival analyses. **a,** Representative image of tumor burden in a mouse brain (sagittal section) bearing orthotopic xenograft of SU-DIPGXIIIpons* xenografted to the pons at endpoint. Survival analysis presented in Figure 4g. White denotes HNA (tumor cells); DAPI nuclei are shown in blue. **b** and **c**, Representative images of tumors at survival endpoint for Figure 4h. **b,** Orthotopic xenograft of SU-DIPG-VI into pons (sagittal section of mouse brain), and in **c,** cortical orthotopic xenograft of SU-pcGBM2 (coronal section of mouse brain). White denotes HNA (tumor cells); Green denotes GFP (tumor cells); DAPI nuclei are shown in blue. **d,** Representative image of mouse brain (sagittal section) from SU-DIPGXIIIpons* xenografted to the pons treated with entrectinib at 120mg/kg PO at endpoint in survival analyses (presented in Figure 4j). White denotes HNA (tumor cells); DAPI nuclei are shown in blue. Scale bar for all = 2000µm.

**Extended Data Fig. 8.**
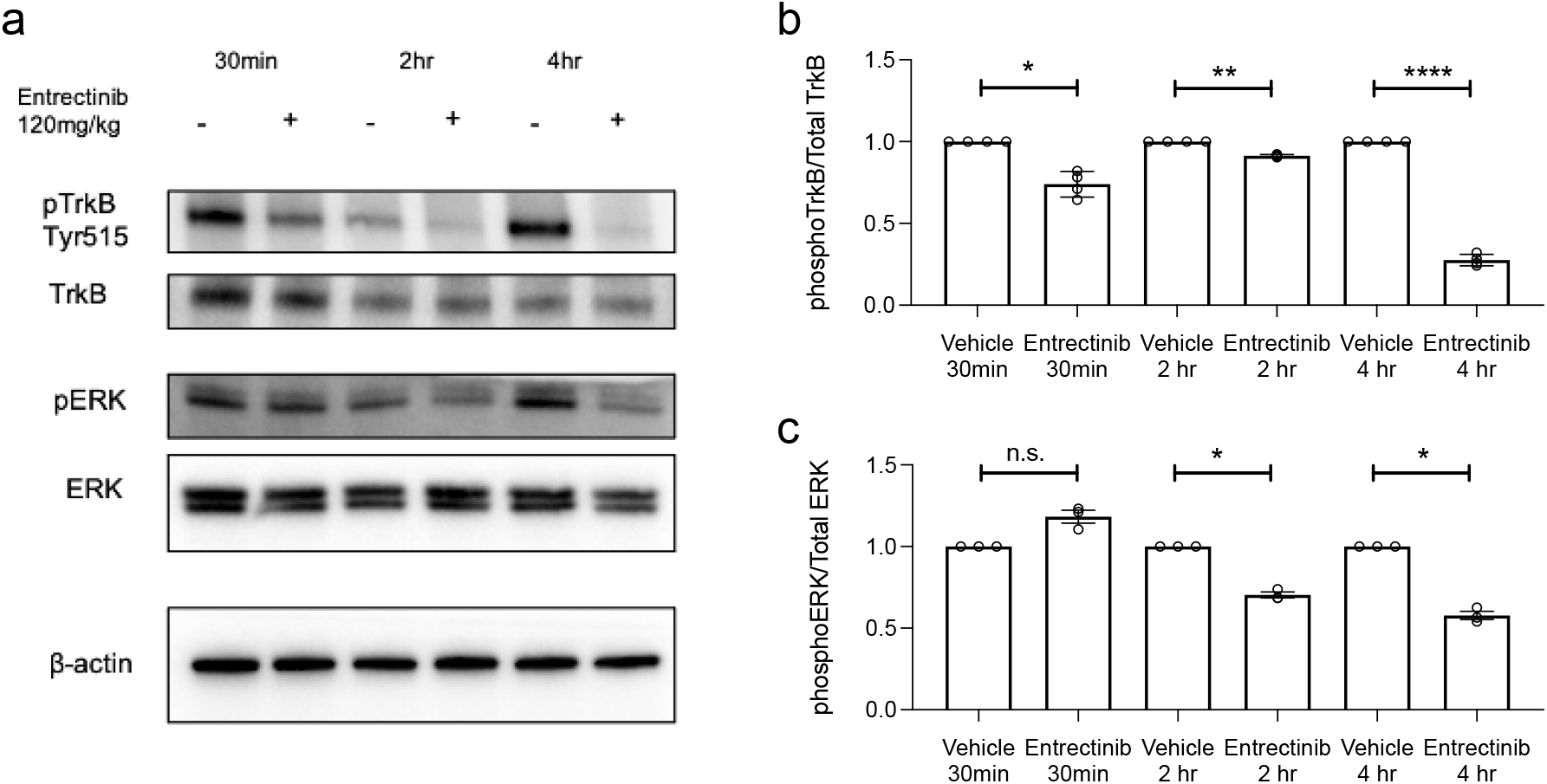
Trk inhibitor brain penetration. **a,** Western blot of whole brain protein lysate collected from NOD-SCID-gamma (NSG) mice that were either treated with one PO dose of 120mg/kg of entrectinib or one dose of vehicle control (PO). The mouse brains were harvested after transcardial perfusion and mice were collected at either 30min, 2 hr and 4 hr post vehicle of entrectinib dosing. The protein lysate was probed for the indicated antibodies to demonstrate inhibition of BDNF-TrkB signaling as an indication of effective drug penetration into brain tissue. **c,** Quantification of TrkB phosphorylation by comparing the ratio of the normalized phospho-TrkB (Tyr515) levels to corresponding total TrkB protein levels between the entrectinib treated and vehicle control mice (*y* axis is in arbitrary units, *n* = 3 technical replicates). **c,** Quantification of MAPK pathway activation by comparing the ratio of the normalized phospho-ERK (T202/Y204) to corresponding total protein levels between the entrectinib treated and vehicle control treated mice (*y* axis is in arbitrary units, *n* = 3 technical replicates). Data are mean ± s.e.m. *P< 0.05, **P< 0.01, ****P<0.0001, one-way analysis of variance (ANOVA) with Tukey’s post hoc analysis.

## Supplementary

**Supplementary Table S1** – Analysis of H3K27M+ DMG biopsy single cell RNASeq data. **a,** Expression level of *NTRK2* and *BDNF* in H3K27M+ DMG. **b,** correlation of genes co-expressed with *NTRK2* in glioma. **c,** GO analysis of top 92 genes co-expressed with *NTRK2* in H3K27M+ DMG. **d,** KEGG analysis of top 92 genes co-expressed with *NTRK2* in H3K27M+ DMG.

**Supplementary Note 1** – Discussion of results of scRNASeq analyses provided in Extended Data Fig.1.

## MATERIALS AND METHODS

### Bioinformatic analysis

For scRNAseq processing, RSEM normalized gene abundances for the Filbin data set were downloaded from the Single Cell Portal (https://singlecell.broadinstitute.org/single_cell, GEO accession: GSE102130). The data were log transformed with a pseudocount of one. The top 50 principal components were calculated using the top 10,000 most variable genes and batch normalized using Harmony until convergence ^51^. The normalized principal components were used to calculate the UMAP embedding. Bootstrapping with 500 iterations was used to calculate 95% confidence intervals. Pathway enrichment analyses were conducted using genes with a bootstrapped lower 95% confidence interval Pearson correlation > 0.25, resulting in a list of 92 genes. The R package clusterProfiler v3.18.1 was used to perform and visualize GO and KEGG enrichment analyses {<u>Yu et al., 2012</u>}. Stemness and lineage signature scores were calculated as described in ^52^. All analysis was performed using the Scanpy Python package ^53^. For synaptic gene co-expression with *NTRK2*, pairwise Pearson correlation for all synaptic genes (gene list as previously published ^4^) was calculated using malignant cells from patient biopsies.

For *NTRK2* and *BDNF* expression levels in DIPG patient-derived cell cultures, FPKM data were analysed from datasets kept in-house and are publicly available to download (GEO accession: GSE94259). Cultures included were SU-DIPG-IV, SU-DIPG-VI, SU-DIPGXIII-p, SU-DIPG-XVII, SU-DIPGXXI, SU-DIPG25, SU-DIPG-XIII-FL).

For TrkB isoform analysis, our previously published ^4^ scRNASeq dataset of patient-derived orthotopic xenograft models was used. NTRK2 isoform abundances were quantified from FASTQ files using Kallisto ^54^. Reads were pseudoaligned against a reference transcriptome created from the hg38 reference genome using Ensembl hg38 transcript annotations. Estimated counts were library-size normalized to counts per million and log transformed with a pseudocount of one. Cells with libraries containing irregular GC content were identified and removed from analysis, resulting in a total of 321 cells.

For BDNF treated tumor cells, RNA was extracted from pelleted cell culture samples using the RNeasy Isolation kit (Qiagen) as per manufacturer’s instructions. Total RNA samples were submitted to Stanford Functional Genomics Facility. RNA integrity was established with Bioanalyzer trace (Agilent). The mRNA was prepared for sequencing using the KAPA Stranded mRNAseq Library prep kit (KK8420), and libraries were indexed with Truseq RNA UD from Illumina (#20021454) as per manufacturer’s instructions. The sequencing was performed on the Illumina NextSeq 500.

Reads were mapped to hg19 annotation using Tophat2 ^55^ (version 2.0.13) and transcript expression was quantified against RefSeq gene annotations using featureCounts ^56^. Differential gene expression and log2 fold change calculations were determined using the DESeq2 package in R ^57^. Volcano plot analysis was conducted using the R-based ‘EnhancedVolcano’ package (https://github.com/kevinblighe/EnhancedVolcano).

### Mice and housing conditions

All animal experiments were conducted in accordance with protocols approved by the Stanford University Institutional Animal Care and Use Committee (IACUC) and performed in accordance with institutional guidelines. Animals were housed according to standard guidelines with unlimited access to water and food, under a 12 hour light : 12 hour dark cycle. For brain tumor xenograft experiments, the IACUC has a limit on indications of morbidity (as opposed to tumor volume). Under no circumstances did any of the experiments exceed the limits indicated and mice were immediately euthanized if they exhibited signs of neurological morbidity or if they lost 15% or more of their initial body weight.

For the Bdnf^TMKI^ mice (C57BL/6J background), knockin mutations in three calcium regulatory element binding sites in the Bdnf promoter IV: CaRE1, CaRE2 and CaRE3/CREB (M. Greenberg: ^44^) were bred to Thy1::ChR2^+/-^ mice (line 18, The Jackson Laboratory, C57BL/6J background) to produce the Bdnf^TMKI^; Thy1::ChR2^+/-^ genotype. These mice were then intercrossed with NSG mice (NOD-SCID-IL2R gamma chain-deficient, The Jackson Laboratory) to produce a Bdnf^TMKI^; Thy1::ChR2^+/-^; NSG genotype to facilitate to facilitate orthotopic xenografting.

### Orthotopic xenografting

For all xenograft studies, NSG MICE (NOD-SCID-IL2R-gamma chain-deficient, The Jackson Laboratory) were used. Male and female mice were used in cohorts equally. For electrophysiological, immunoelectron microscopy and calcium imaging experiments, a single cell suspension from SU-DIPGVI and SU-DIPGXIIIFL were injected into the hippocampal region. For survival analysis, DIPG cultures (SU-DIPGVI, SU-DIPGXIIIp*) were injected into the pontine region and for cortical GBM cultures (SU-pcGBM2), cells were injected into the cortex. A single cell suspension of all cultures were prepared in sterile culture medium (see *cell culture*) immediately before surgery. Animals at postnatal day (P) 28-35 were anaesthetized with 1-4% isoflurane and placed on stereotactic apparatus. Under sterile conditions, the cranium was exposed via a midline incision and a 31-gauge burr hole made at exact coordinates. For hippocampal injections the coordinates were as follows: 1.5mm lateral to midline, 1.8mm posterior to bregma, −1.4 deep to cranial surface. For pontine injections coordinates were: −1mm lateral to midline, 0.8mm posterior to lambda, −5 deep to cranial surface. For cortical injections coordinates were: 1mm anterior to bregma, 0.5mm lateral to midline, 1.7mm deep to cranial surface. Cells were injected using a 31-gauge Hamilton syringe at an infusion rate of 0.4 µl min^-1^ with a digital pump. At completion of infusion, the syringe needle was allowed to remain in place for a minimum of 2min, then manually withdrawn. The wound was closed using 3M Vetbond (Thermo Fisher) and treated with Neo-Predef with Tetracaine Powder.

### Survival studies

For survival studies, mice were xenografted at 4-5 weeks of age with the cultures SU-DIPG-VI (WT and NTRK2 KO), SU-pcGBM2 (WT and NTRK2 KO) and SU-DIPGXIIIp*. After xenografts, mice were continuously monitored for signs of neurological deficits or health decline. For inhibitor treatment, SU-DIPGXIIIp* was treated with 120mg/kg PO daily of entrectinib (HY-12678, MedChemExpress, 7% DMSO (Sigma), 10% Tween 80 (Sigma) in sterile H2O) for 14 days, starting at 2 weeks post xenograft. Morbidity criteria were a 15% reduction in weight or severe neurological motor deficits consistent with brain dysfunction (brainstem tumors exhibited circling and barrel roles, cortical tumors displayed seizures and loss of gait). Statistical analysis were performed with Kaplan-Meier survival analysis using log rank testing.

### Slice preparation for electrophysiology and calcium imaging experiments

Coronal slices (300 µm thick) containing the hippocampal region were prepared from mice (4-8 weeks after xenografting) in accordance with a protocol approved by Stanford University APLAC. After rapid decapitation, the brain was removed from the skull and immersed in ice-cold slicing artificial cerebrospinal fluid (ACSF) containing (in mM): 125 NaCl, 2.5 KCl, 25 glucose, 25 NaHCO_3_ and 1.25 NaH_2_PO_4_, 3 MgCl_2_ and 0.1 CaCl_2_. After cutting, slices were incubated for 30 min in warm (30 °C) oxygenated (95% O_2_, 5% CO_2_) recovery ACSF containing (in mM): 100 NaCl, 2.5 KCl, 25 glucose, 25 NaHCO_3_, 1.25 NaH_2_PO_4_, 30 sucrose, 2 MgCl_2_ and 1 CaCl_2_ before being allowed to equilibrate at room temperature for an additional 30 min.

### Electrophysiology

Slices were transferred to a recording chamber and perfused with oxygenated, warmed (28–30 °C) recording ACSF containing (in mM): 125 NaCl, 2.5 KCl, 25 glucose, 25 NaHCO3, 1.25 NaH2PO4, 1 MgCl2 and 2 CaCl2. Tetrodotoxin (0.5 μM) was perfused with the recording ACSF to prevent neuronal action potential firing. Slices were visualized using a microscope equipped with DIC optics (Olympus BX51WI). Recording patch pipettes (2–3 MΩ) were filled with K-gluconate-based pipette solution containing (in mM): 130 K-gluconate, 20 KCl, 5 Na2-phosphocreatine, 10 HEPES, 4 Mg-ATP, 0.3 GTP, and 50 μM Fluo-4, pH = 7.3. Pipette solution additionally contained Alexa 568 (50 μM) to visualize the cell through dye-filling during whole-cell recordings. Glutamate (1 mM) in recording ACSF was applied via a puff pipette approximately 100 μm away from the patched cell. Recombinant BDNF human protein (Peprotech, #450-02) was added to ACSF at 100ng/ml, in addition to tetrodotoxin (0.5 μM) for perfusion. Signals were acquired with a MultiClamp 700B amplifier (Molecular Devices) and digitized at 10 kHz with an InstruTECH LIH 8+8 data acquisition device (HEKA). Data were recorded and analyzed using AxoGraph X (AxoGraph Scientific) and IGOR Pro 8 (Wavemetrics).

### Calcium imaging

SU-DIPGVI and SU-DIPG-XIII-FL were transduced with lentivirus containing the genetically encoded calcium indicator GCaMP6s (pLV-ef1-GCAMP6s-P2A-nls-tdTomato) as published previously ^4^. Cells were xenograft into the CA1 region of the hippocampus as described above.

Calcium imaging experiments were performed on *in situ* xenograft slices were visualized using a microscope equipped with DIC optics (Olympus BX51WI). Slices were perfused with oxygenated aCSF, as described above, at a constant temperature of (28–30 °C) and containing (in mM): 125 NaCl, 2.5 KCl, 25 glucose, 25 NaHCO3, 1.25 NaH2PO4, 1 MgCl2 and 2 CaCl2. Tetrodotoxin (0.5 μM) was perfused with the recording ACSF to prevent neuronal action potential firing. Glutamate (1 mM) in recording ACSF was applied via a puff pipette (250msec) approximately 100 μm away from the tDTomato expressing cells. Recombinant BDNF human protein (Peprotech, #450-02) was added to ACSF at 100ng/ml, in addition to tetrodotoxin (0.5 μM) for perfusion. Excitation light was at 594 (for TdTomato) and 488 (for GCaMP6s) provided by pE-300^ULTRA^ (CoolLED). The recording software used was FlyCapture2 (Point Grey) and analyzed using ImageJ.

### Cell culture

Primary patient derived cultures were derived as described previously ^1 58^ with informed consent and Institutional Review Board (IRB) approval granted. All cultures are monitored by short tandem repeat (STR) fingerprinting for authenticity throughout the culture period and mycoplasma testing was routinely performed.

Cultures are grown as neurospheres (unless otherwise stated) in serum free medium consisting of DMEM (Invitrogen, Carlsbad, CA), Neurobasal(-A) (Invitrogen, Carlsbad, CA), B27(-A) (Invitrogen, Carlsbad, CA), heparin (2ng/mL), human-bFGF (20ng/mL) (Shenandoah, Biotech, Warwick, PA), human-bEGF (20ng/mL) (Shenandoah, Biotech, Warwick, PA), human-PDGF-AA (10ng/mL) (Shenandoah, Biotech, Warwick, PA), human-PDGF-BB (10ng/mL) (Shenandoah, Biotech, Warwick, PA). The spheres were dissociated using TrypLE (Gibco) for seeding of *in vitro* experiments.

### Biotinylation

Glioma cells (SU-DIPG-VI) were seeded on laminin coated wells of 6-well plates at a density of 500,000 cells. One day after plating, the medium was changed to medium without growth factors to “starve” the cells for three days. Cells were treated with either vehicle (equal volume added of 0.1% BSA in H2O) or 100nM BDNF recombinant protein (Peprotech, #450-02, stock 0.25µg/µl in 0.1% BSA in H2O) for specified time periods. To label surface proteins, the cells were washed twice with ice cold PBS befpre adding 1mg/mL sulfo-NHS-SS-biotin (Thermo Scientific) for 10min at 4°C with continuous gentle shaking. The reaction was quenched (100mM glycine, 25mM Tris-HCL, pH 7.4) for 5min and then washed in ice-cold PBS three times; all procedures were carried out at 4°C. The biotinylated cells were then lysed in RIPA lysis buffer (Santa Cruz Biotechnology) supplemented with PMSF, protease inhibitor cocktail and sodium orthovanadate as per manufacturers recommendations. Insoluble material was removed by centrifugation at 10,000xg at 4°C for 10min and the supernatant was incubated with 50ul NeutrAvidin agarose resin (Thermo Scientific) with gentle mixing overnight at 4°C. Beads were washed with lysis buffer three times and proteins bound to the beads were eluted with NuPage LDS and sample reducing buffer (Life Technologies).

### Western blot analysis

For ligand activation experiments, patient-derived cultures were incubated in media supplemented with only B-27 supplement minus vitamin A for three days. Cells were dissociated using 5mM EDTA in HBSS and resuspended in media with B-27 supplement for 4 hours before incubation with recombinant BDNF protein (100nM, Peprotech, #450-02) or vehicle (equal volume of 0.1% BSA in H2O). After ligand stimulation for the stated time-points the cells were washed in PBS, before lysis using the RIPA Lysis Buffer System containing PMSF, protease inhibitor cocktail and sodium orthovanadate (Santa Cruz Biotechnology). Following quantification using the Pierce BCA Protein Assay Kit (Thermo Fisher Scientific), equal amounts of total protein were loaded onto for each sample for standard western blot.

### Mice and housing conditions

All experiments were performed in accordance with Stanford University Institutional Animal Care and Use Committee (IACUC) approved protocols and conducted using institutional guidelines. Animal housing followed standard guidelines, with free access to water and food in a 12h light:12h dark cycle. Mice were continuously monitored for their condition, as per IACUC guidelines, and euthanized upon signs of morbidity (neurological symptoms or a loss of >15% body weight).

### Neuron-glioma co-culture

For synaptic puncta assays, neurons were isolated from CD1 (The Jackson Laboratory) at P_1_ using he Neural Tissue Dissociation Kit – Postnatal Neurons (Miltenyi), and followed by the Neuron Isolation Kit, Mouse (Miltenyi) per manufacturers instructions. After isolation 200,000 neurons were plated onto circular glass coverslips (Electron Microscopy Services) pre-coated with poly-L-lysine (Sigma) and mouse laminin (Thermo Fisher) as described previously (Venkatesh). Neurons were cultured in BrainPhys neuronal medium containing B27 (Invitrogen), BDNF (10ng ml^-1^, Shenandoah), GDNF (5ng ml^-1^, Shenandoah), TRO19622 (5µM; Tocris), β-mercaptoethanol (Gibco). The medium was replenished on days in vitro (DIV) 1. On DIV 5, fresh media was added containing 50,000 glioma cells expressing PSD95-RFP. The neuron-glioma culture was incubated for 72hours and then fixed with 4% paraformaldehyde (PFA) for 20min at room temperature and stained for immunofluorescence analysis.

### Cloning constructs

For SEP-GluA2(Q)-TagBFP, addgene plasmid EFS-Cas9-Puro (#138317) was digested with AgeI, MluI, and EcoRV; the 6kb lentiviral backbone was isolated via gel extraction. The SEP fragment with Gibson overhangs was amplified from Addgene plasmid pCI-SEP-GluR2(Q) (#24002) with primers: 5’-AACGGGTTTGCCGCCAGAACACAGGACCGGTGCCACCATGCAAAAGATTATGCAT ATTTC and 5’-CCCCCTATCTGTATGCTGTTGCTAGCTTTGTATAGTTCATC. Human *GRIA2* (GluA2) with Gibson overhangs was amplified from pLV-EF1a-GFP-GRIA2 (described in Venkatesh et al., 2019) in two parts to introduce R583Q. GRIA-part 1 was amplified with primers: 5’-CAAAGCTAGCAACAGCATACAGATAGGG and 5’-TTGGCGAAATATCGCATCCCTGCTGCATAAAGGCACCCAAGGA. GRIA-part 2 was amplified with primers: 5’-TTATGCAGCAGGGATGCGATATTTCGCCAA and 5’-TCTTCGACATCTCCGGCTTGTTTCAGCAGAGAGAAGTTTGTTGCGCCGGATCCAATTTTAACACTTTCGATGC. TagBFP2 with Gibson overhangs was amplified from Addgene plasmid pLenti6.2-TagBFP (#113724) with primers: 5’-TCTGCTGAAACAAGCCGGAGATGTCGAAGAGAATCCTGGACCGATGAGCGAGCTGATTAAG and 5’-TTGTAATCCAGAGGTTGATTGTCGACTTAACGCGTTTAATTAAGCTTGTGCCC. DNA fragments above were stitched together using Gibson Assembly and transformed.

For PSD95-PURO, Addgene plasmid EFS-Cas9-Puro (#138317) was digested with AgeI, BamHI, and EcoRV; the 6.7kb lentiviral backbone was isolated via gel extraction. PSD95-RFP with Gibson overhangs was amplified from PSD95-RFP (described in Venkatesh et al., 2019) with primers: 5’-TCGCAACGGGTTTGCCGCCAGAACACAGGTCTAGAGCCACCATGGACTGTCTCTG TATAG and 5’-TGTTTCAGCAGAGAGAAGTTTGTTGCGCCGGATCCATTAAGTTTGTGCCCCAG. DNA fragments above were stitched together using Gibson Assembly and transformed.

### CRISPR deletion

Target sequencing for sgRNA was generated using the online predictor at https://cctop.cos.uni-heidelburg.de. The validated sgRNA sequence used for *NTRK2* deletion in all cultures was 5’-GTCGCTGCACCAGATCCGAG – 3’. The scrambled control sequence used was 5’-GGAGACGTGACCGTCTCT −3’. The custom oligos were purchased from Elim Biopharmaceuticals. The oligos were phosphorylated in a reaction with the oligo (10uM), 1x T4 DNA ligase buffer (B0202, NEB) and T4 PNK (M020, NEB) with a program 45min 37°C, 2min30sec at 95°C, cool 0.1°C/sec to 22°C. The sgRNA was cloned into the Lenti vector (pL-CRISPR.EFS.RFP, Addgene #57819). First the vector was digested in a reaction with Fast Digest Buffer (B64, Thermo fisher), BsmBI restriction enzyme (FD0454, Thermo Fisher), DTT (10mM) with program 45min at 37°C, heat inactivate 10min at 65°C. The digested vector backbone was dephosphorylated using Antartica phosphatase (M0289, NEB) in Antarctic phosphatase buffer (B0289, NEB) at 37°C for 30min, before purifying after running on a 1% agarose gel. The phosphorylated oligo duplexes were ligated into the vector backbone in a reaction with T4 DNA ligase buffer and T4 DNA ligase and incubated at room temperature for 1 hour. Stabl3 (Invitrogen) cells were transformed with the assembled plasmids and individual colonies picked the next day for propagation and sanger sequencing (ElimBio). Lentiviral particles for were produced following transfection of the lentiviral packaging vectors (pΔ8.9 and VSV-g) and either the *NTRK2* CRISPR vector or the control scramble vector into HEK293T cells and collected 48hours later. The viral particles were concentrated using Lenti-X Concentrator (Takara Bio) and resuspended in TSM base and stored at −80°C for future use. the RFP positive cells were FACs sorted for purity and returned to culture. Lentiviral particles for shRNA knockdown of *NTRK2* were produced following transfection of the lentiviral packaging vectors (pΔ8.9 and VSV-g) and either the *PIK3CA* shRNA vector (TRCN0000197207; Sigma) or the control shRNA vector (SHCOO2; Sigma) into HEK293T cells and collected 48hours later. The viral particles were concentrated using Lenti-X Concentrator and resuspended in TSM base and stored at −80°C for future use. Control or *PIK3CA* shRNA lentiviral particles were transduced into SU-DIPGVI cultures and the transduced cells were selected with puromycin (4 μg/mL) from day 3.

### pHluorin live imaging

Glioma cells (SU-DIPG-VI) expressing the SEP-GluA2(Q)-TagBFP and PSD95-RFP-Puro constructs (see *Cloning constructs*) were cultured as adherent cells, with mouse neurons, on laminin coated 27mm glass bottom plates (#150682, Thermo Scientific). ACSF was made at pH 7.4 (See *Electrophysiology*) and at pH 5.5 using the membrane impermeable acid, MES hydrate (Sigma) to replace NaHCO_3_ at equimolar concentration. The ACSF was perfused onto the culture dish using a 3D-printed custom-built stage and tubing for manual perfusion of the solution. Images were collected using a Zeiss LSM980 confocal microscope equipped with a plexiglass environmental chamber, heated stage and CO2 module, and post-processed with Airyscan. The cells were kept at 37°C with 5% CO2 for the duration of the imaging period. Puncta were identified as bright spots, with colocalisation of the PSD95-RFP signal to the GFP signal from the SEP-GluA2(Q) puncta. In ImageJ, an ROI was manually drawn over the PSD95-RFP signal and used to measure both the RFP and SEP signal, thus blinding the area chosen for the SEP-GluA2(Q) signal. All puncta were identified for the first timepoint and followed for all the subsequent time images, thus the choice was blind with respect to outcome. The levels of SEP-GluA2(Q) were represented as a ratio to the levels of PSD95-RFP, to account for any fluorescence intensity changes that may occur due to photobleaching, or Z-Axis drifting, during the imaging time course. For BDNF perfusion experiments, ACSF (pH 7.4) containing 100nM BDNF (Peprotech, #450-02) was perfused into the chamber. After imaging the signal in response to BDNF, the signal was then quenched with pH 5.5 to confirm the puncta of interest were membrane bound GluA subunits. The fluorescence intensity was measured using ImageJ.

### Immuno-electron microscopy

Twelve weeks post xenografting, mice were euthanized by transcardial perfusion with Karnovsky’s fixative: 4% paraformaldehyde (EMS 15700) in 0.1M sodium cacodylate (EMS 12300), 2% glutaraldehyde (EMS 16000), p.H 7.4. For all xenograft analysis, transmission electron microscopy was performed in the tumor mass located in the CA1 region of the hippocampus. At room temperature the samples were post fixed in 1% osmium tetroxide (EMS 19100) for 1 hour, washed 3 times with ultrafiltered water, before 2-hour en bloc staining. The samples were dehydrated in graded ethanol (50%, 75% and 95%) for 15 min each at 4°C before equilibrating to room temperature and washed in 100% ethanol twice, followed by a 15 min acetonitrile wash. Samples were immersed for 2 hours in Embed-812 resin (EMS 14120) with 1:1 ratio of acetonitrile, followed by a 2:1 Embed-812:acetonitrile for 2 hours, then in Embed-812 for 2 hour. The samples were moved to TAAB capsules with fresh resin and kept at 65°C overnight. Sections of 40 and 60 nm were cut on an Ultracut S (Leica) and mounted on 100-mes Ni grids (EMS FCF100-Ni). For immunohistochemistry, microetching was done with 10% periodic acid and eluting of osmium with 10% sodium metaperiodate for15 min at room temperature on parafilm. Grids were rinsed with water three times, followed by 0.5 M glycine quench, and then incubated in blocking solution (0.5% BSA, 0.5% ovalbumin in PBST) at room temperature for 20 min. Primary rabbit anti-GFP (1:300; MBL International) was diluted in the same blocking solution and incubated overnight at 4 °C. The next day, grids were rinsed in PBS three times, and incubated in secondary antibody (1:10 10-nm gold-conjugated IgG TED Pella15732) for one hour at room temperature and rinsed with PBST followed by water. For each staining set, samples that did not contain any GFP-expressing cells were stained simultaneously to control for any non-specific binding. Grids were contrast stained for 30 s in 3.5% uranyl acetate in 50% acetone followed by staining in 0.2% lead citrate for 90 s. Samples were imaged using a JEOL JEM-1400 TEM at 120 kV and images were collected using a Gatan Orius digital camera.

### Conditioned media

Wild-type or TMKI mice with Thy1::Chr2 were used at 4-7 weeks of age. Brief exposure to isoflurane rendered the mice unconscious before immediate decapitation. Extracted brains were placed in an oxygenated sucrose cutting solution containing xxx and sliced at 350um as described previously (Venkatesh 2015). The slices were placed in ACSF (see *Electrophysiology*) and allowed to recover for 30 minutes at 37C and 30mins at room temperature. After recovery the slices were moved to fresh ACSF and stimulated using a blue-light LED using a microscope objective. The stimulation paradigm was 20-Hz pulses of blue light for 30 seconds on, 90 seconds off over a period of 30 minutes. The surrounding conditioned medium was collected and used immediately or frozen at −80°C for future use.

### Immunohistochemistry

Animals were anaesthetized with intraperitoneal avertin (tribromoethanol), then transcardially perfused with 20 ml of PBS. Brains were fixed in 4% PFA overnight at 4 °C, then transferred to 30% sucrose for cryoprotection. Brains were then embedded in Tissue-Tek O.C.T. (Sakura) and sectioned in the coronal plane at 40 μm using a sliding microtome (AO 860, American Optical).

For immunohistochemistry, coronal or sagittal sections were incubated in blocking solution (3% normal donkey serum, 0.3% Triton X-100 in TBS) at room temperature for 30 min. Mouse anti-human nuclei clone 235-1 (1:250; Millipore), rabbit anti-histone H3.3K27M mutant (1:500; abcam) were diluted in antibody diluent solution (1% normal donkey serum in 0.3% Triton X-100 in TBS) and incubated overnight at 4 °C. Sections were then rinsed once with TBS, before an incubation with DAPI (1µg/ml in TBS, Thermo Fisher) and then another rinse with TBS. Slices were incubated in secondary antibody solution; Alexa 594 donkey anti-rabbit IgG, Alexa 647 donkey anti-mouse IgG, all used at 1:500 (Jackson Immuno Research) in antibody diluent at 4 °C overnight. Sections were washed three times with TBS and mounted with ProLong Gold Mounting medium (Life Technologies).

### EdU incorporation assay

EdU staining was performed on *in vitro* cell culture slides or on glass coverslips in 24-well plates which were precoated with poly-l-lysine (Sigma) and laminin (Thermo Fisher). Neurosphere culture were dissociated with TrypLE and plated onto coated slides, once the cells had adhered the media was replaced with growth factor-depleted media for 72hours. Recombinant protein (BDNF, peprotech), inhibitors (entrectinib HY-12678 and Larotrectinib HY-12866, both MedChem Express) and vehicle (0.1% BSA and/or DMSO) were added for specified times with 10uM of EdU. After added 24 hours the cells were fixated with 4% paraformaldehyde in PBS for 20mins and then stained using the Click-iT EdU kit and protocol (Invitrogen, Carlsbad, CA) and mounted using Prolong Gold mounting medium with DAPI (Life Technologies). Proliferation index was determined by quantifying the fraction of EdU labeled cells/DAPI labeled cells using confocal microscopy at 20x magnification.

### Statistical analyses

Statistical tests were conducted using Prism v9.1.0 (GraphPad) software unless otherwise indicated. Gaussian distribution was confirmed by the Shapiro-Wilk normality test. For parametric data, unpaired two-tailed Student’s t-test or one-way ANOVA with Tukey’s post hoc tests to examine pairwise differences were used as indicated. Paired two-tailed Student’s t-tests were used in the case of same cell experiments (as in electrophysiological recordings). For data normalized to a control mean (as in western blot or pHluorin analyses) One Sample *t*-test were used against the mean of the control (either 0 or 1), with Wilcoxon signed-rank test for non-parametric data. For non-parametric data, a two-sided unpaired Mann-Whitney test was used as indicated, or a one-tailed Wilcoxon matched pairs signed rank test was used for same-cell experiments. Two-tailed log rank analyses were used to analyse statistical significance of Kaplan-Meier survival curves. On the basis of variance of xenograft growth in control mice, we used at least three mice per genotype to give 80% power to detect effect size of 20% with a significance level of 0.05. For all mouse experiments, the number of independent mice used is listed in the figure legend.

## Data availability

RNA sequencing of patient-derived cultures treated with recombinant BDNF protein will be made available on Gene Expression Omnibus (GEO) prior to publication. All other data are available in the manuscript or from the corresponding author upon reasonable request. Source data will be uploaded with the final version of the manuscript.

## Code availability

Sources for all code used have been provided, no custom code was created for this manuscript.

## ACKNOWLEDGMENTS

This work was supported by grants from the National Institute of Neurological Disorders and Stroke (R01NS092597 to M.M.), NIH Director’s Pioneer Award (DP1NS111132 to M.M.), National Cancer Institute (P50CA165962), Abbie’s Army (to M.M.), Robert J. Kleberg, Jr. and Helen C. Kleberg Foundation (to M.M.), Cancer Research UK (to M.M.), Damon Runyon Cancer Research (to K.R.T.), N8 Foundation (to M.M.), McKenna Claire Foundation (to M.M.), Kyle O’Connell Foundation (to M.M.), Virginia and D.K. Ludwig Fund for Cancer Research (to M.M.), Waxman Family Research Fund (to M.M.), Will Irwin Research Fund (to M.M.)

## Competing Interests

M.M. is on the SAB for Cygnal Therapeutics.

## AUTHOR CONTRIBUTIONS

K.R.T and M.M. designed, conducted, and analysed experiments. T.B. conducted electrophysiology experiments. K.R.T. conducted calcium imaging experiments, xenografting for all experiments, pHluorin confocal imaging, in vitro and in vivo data collection and analyses. L.N. and K.R.T contributed to electron microscopy data acquisition, and K.R.T and H.S.V. to analyses. P.D. performed single-cell transcriptomic analyses. R.M. performed RNASeq data analyses. K.R.T and H.Z contributed to synaptic puncta confocal imaging. K.R.T performed western blot analyses. G.H. generated constructs. A.P. performed CRISPR deletion. A.H., B.Y. and I.C. contributed to in vivo data collection. K.R.T., H.S.V., T.B., G.H., and M.M. contributed to manuscript editing. K.R.T. and M.M. wrote the manuscript. M.M. conceived of the project and supervised all aspects of the work.

## Notes

### Competing Interest Statement

M.M. is on the SAB of Cygnal Therapeutics

